# A Multi-task Platform for Profiling Cognitive and Motivational Constructs in Humans and Nonhuman Primates

**DOI:** 10.1101/2023.11.09.566422

**Authors:** Marcus R. Watson, Nathan Traczewski, Seema Dunghana, Kianoush Banaie Boroujeni, Adam Neumann, Xuan Wen, Thilo Womelsdorf

**Author notes:** Corresponding Authors: Dr. Thilo Womelsdorf, Vanderbilt University, Psychology Department, 301 Wilson Hall, 111 21st Avenue South, 37240-1103 Nashville TN.

## Abstract

**Background:** Understanding the neurobiological substrates of psychiatric disorders requires comprehensive evaluations of cognitive and motivational functions in preclinical research settings. The translational validity of such evaluations will be supported by (1) tasks with high construct validity that are engaging and easy to teach to human and nonhuman participants, (2) software that enables efficient switching between multiple tasks in single sessions, (3) software that supports tasks across a broad range of physical experimental setups, and (4) by platform architectures that are easily extendable and customizable to encourage future optimization and development.

**New Method:** We describe the *Multi-task Universal Suite for Experiments* (*M-USE*), a software platform designed to meet these requirements. It leverages the Unity video game engine and C# programming language to (1) support immersive and engaging tasks for humans and nonhuman primates, (2) allow experimenters or participants to switch between multiple tasks within-session, (3) generate builds that function across computers, tablets, and websites, and (4) is freely available online with documentation and tutorials for users and developers. M-USE includes a task library with seven pre-existing tasks assessing cognitive and motivational constructs of perception, attention, working memory, cognitive flexibility, motivational and affective self-control, relational long-term memory, and visuo-spatial problem solving.

**Results:** M-USE was used to test NHPs on up to six tasks per session, all available as part of the Task Library, and to extract performance metrics for all major cognitive and motivational constructs spanning the Research Domain Criteria (RDoC) of the National Institutes of Mental Health.

**Comparison with Existing Methods:** Other experiment design and control systems exist, but do not provide the full range of features available in M-USE, including a pre-existing task library for cross-species assessments; the ability to switch seamlessly between tasks in individual sessions; cross-platform build capabilities; license-free availability; and its leveraging of video-engine capabilities used to gamify tasks.

**Conclusions:** The new multi-task platform facilitates cross-species translational research for understanding the neurobiological substrates of higher cognitive and motivational functions.

## 1. Introduction

Preclinical research aims to understand how neurophysiological, genetic, molecular, or cellular processes affect specific cognitive functions. Translating preclinical insights to clinical/neuropsychiatric contexts, and developing and evaluating treatments for specific disorders, requires assessing cognitive functions with high validity across human and animal models (Kangas & Bergman, 2017; Redish et al., 2022). Research with nonhuman primates (NHPs) is an essential component of this research (Oikonomidis et al., 2017; Palmer et al., 2021; Roelfsema & Treue, 2014; Scott & Bourne, 2022), because of the anatomical and functional similarities between human and NHP prefrontal cortex and its associated networks (Passingham & Wise, 2012), which are implicated in many neuropsychiatric disorders..

Previous research has documented that the prefrontal cortex functions with the highest diagnostic value for neuropsychiatry include attention, cognitive flexibility, motivation, affective valuation, problem solving, and relational long-term memory (Friedman & Robbins, 2022; Passingham, 2021). Existing task paradigms evaluate each of these in NHPs and humans (Calapai et al., 2017; Friedman & Robbins, 2022; Oikonomidis et al., 2017; Perdue et al., 2018; Weed et al., 1999), providing domain-specific insights into the constructs that the National Institutes of Mental Health has summarized in their Research Domain Criteria (RDoC) as being central to advance our understanding of the neural correlates of neuropsychiatric disorders (Aragona, 2014; Cuthbert & Insel, 2013; Morris & Cuthbert, 2012).

Assessing such functions across species has been spearheaded by the Cambridge Automated Neuropsychological Test Associated Battery (CANTAB, cf. Fray et al., 1996; Sahakian & Owen, 1992), that includes classical diagnostic tasks used for humans (Langley et al., 2023) and NHPs (Monkey CANTAB, Lafayette Instrument Company, Lafayette, IN). These cross-species assessments have been instrumental for optimizing diagnostic testing, evaluating treatments, and advancing the pre-clinical understanding of neuropsychiatric disorders (Friedman & Robbins, 2022; Palmer et al., 2021). However, while it is routine to assess multiple RDoC constructs in humans using test batteries like CANTAB (Harvey, 2023; Langley et al., 2023), the MATRICS Consensus Cognitive Test Battery (MCCB), CogState (Buchanan et al., 2011; Harvey, 2023), or CNTRICS (Moore et al., 2013), in NHP the assessment of multiple cognitive domains has remained a major challenge with the majority of studies reporting results from one or few tests, and even fewer studies that allow automatized task switching during the assessment (Berger et al., 2018; Calapai et al., 2017; Hassani et al., 2021; Hassani et al., 2023; Moore et al., 2003; Moore et al., 2005; Taffe et al., 2002; Weed et al., 1999; Womelsdorf et al., 2021).

Here, we address this challenge by introducing a novel cross-species, multi-task software platform, the *Multi-Task Universal Suite for Experiments* (M-USE, **Figure 1A**). M-USE has been designed for efficient assessment of RDoC domains using multiple tasks whose performance causally depend on different subfields of primate prefrontal cortex (Passingham, 2021). M-USE’s *Task Library* (**Figure 1A**) contains pre-configured tasks assessing these domains, the platform’s *Experimenter Manual* outlines how to use these tasks for assessment, and the *Developer Manual* outlines how to customize and extend the M-USE platform with novel tasks (see Appendix for all links).

**Figure 1.**
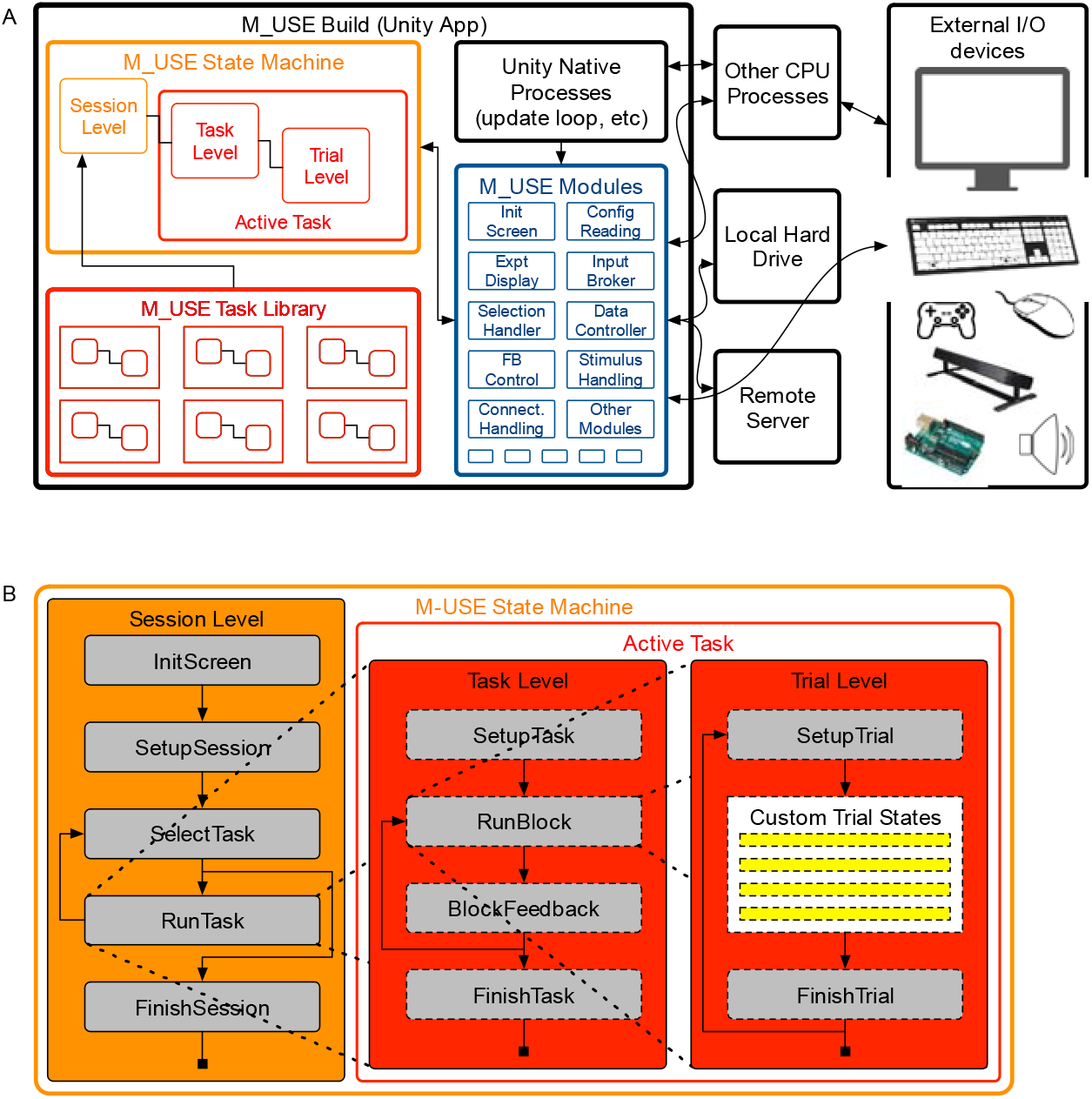
Build architecture and state system of the Multi-task Unified Suite for Experiments (M-USE). (**A**) M-USE builds consist of Unity’s native processes (in black), the M_USE State Machine that controls task operations (orange), a Task Library with pre-configured tasks that can be accessed in a plug-and-play manner by the State Machine (red), and custom Modules that govern I/O and other common experimental needs (blue). All interactions with other CPU processes or experimental equipment are mediated through the Modules, allowing the State Machine and Task Library to remain encapsulated. (**B**) The primary components of the M-USE State Machine include sequences of states at the Session, Task and Trial levels. A developer’s primary work consists in defining the custom Trial states that control their task (in yellow).

M-USE enables direct comparisons of cognitive and motivational performance across species and participant groups because it supports running tasks in multiple contexts, including (*i*) touchscreen setups common in cage-mounted training stations in NHP research, (*ii*) tablet (iPad) devices for convenient testing of humans; (*iii*) classical computer set-ups common to research labs; and (*iv*) online Web Graphics Library (WebGL) applications that enable online internet collection of human data.

We introduce M-USE for users and developers by first reviewing the hierarchical software architecture that enables multi-task handling and the tasks in our Task Library, and then presenting results illustrating the general functioning of the platform, and the specific cognitive and motivational metrics associated with each task.

## 2. Methods

### 2.1 The M-USE Build

M-USE is a custom software package for the Unity video-game engine, comprised of C# scripts and Unity scenes. The executable application is available in the form of *builds* for Windows and Macintosh systems, as well as for WebGL apps hosted on webpages. M-USE builds consist of four distinct, interacting components (**Figure 1A**): Unity’s native processes, the *M-USE State Machine*, *Task Library*, and *Modules*. The state machine is, in formal terms, a *multi-level hierarchical finite state machine* (Wagner et al., 2006), with an always-active Session level that governs access to all lower levels, which included the paired Task and Trial levels that define each specific task stored in the *M-USE Task Library*, (**Figure 1A**). This structure allows tasks to be flexibly accessed by the Session Level as needed (**Figure 1B**), and once one task is run the SelectTask state can be re-started, and different tasks, or the same task with different configurations, can be accessed from the Task Library and run in the same session.

The state machine is entirely encapsulated, with all interactions with other Unity or CPU processes governed by the various M-USE modules (**Figure 1A**). Figure 2 highlights several of these interactions, which we discuss in the following sections.

**Figure 2.**
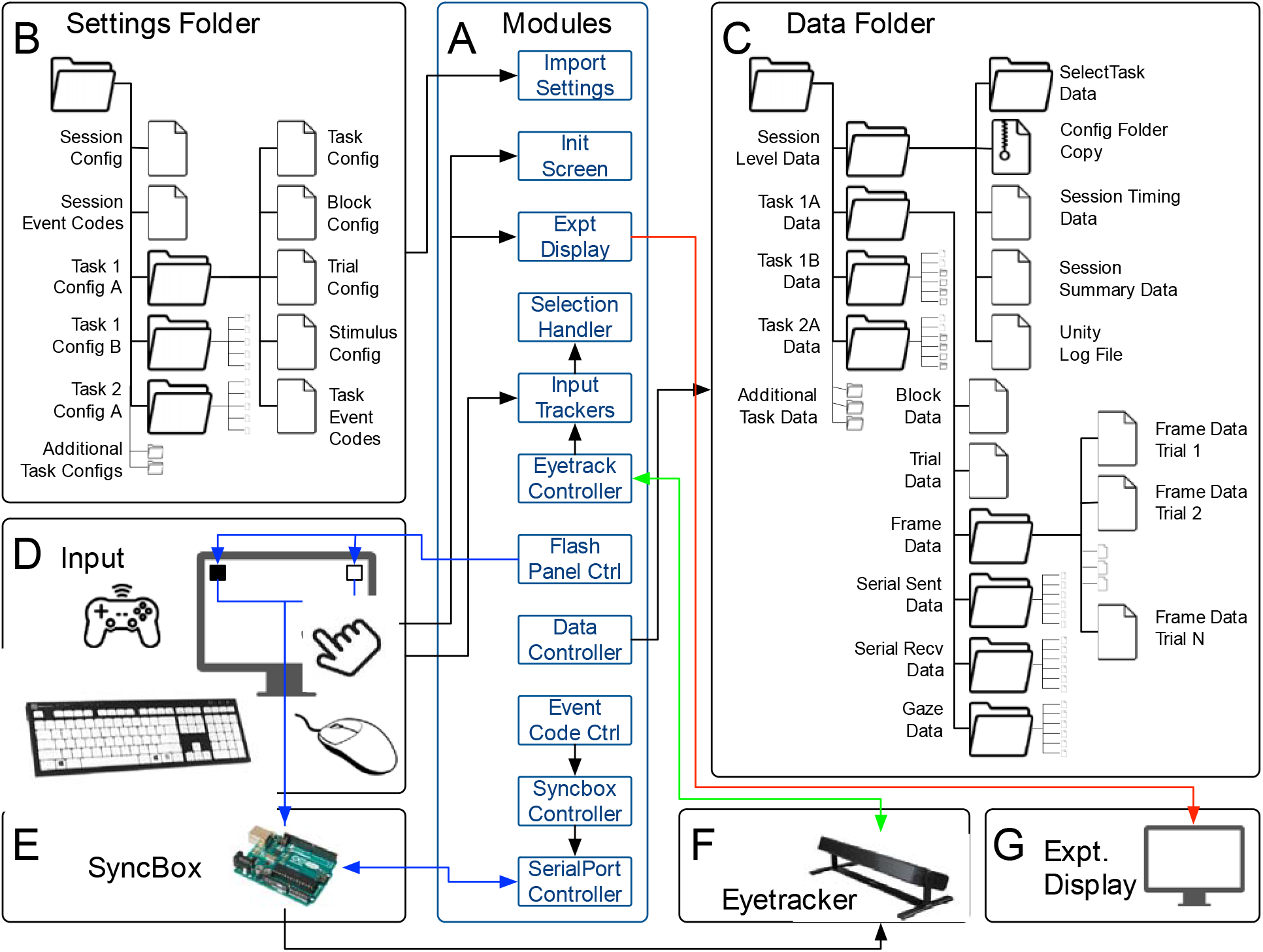
Basic I/O functions of M-USE. (A) Modules control all aspects of I/O, including (B) reading configuration files that apply across a session, or to individual tasks, (C) writing data files for the session and task, (D) receiving participant input, (E) communicating to and from time synchronization devices such as our Arduino-based SyncBox, (F) communicating with eyetrackers, if used, and (G) displaying up to date information and allowing experimenter to manipulate trial variables on an Experimenter Display.

### 2.2 Initialization screen

M-USE sessions begin with an initialization screen with fields to specify participant ID and age, settings and data folder paths, and server details if applicable. These values are retained from the previous session from session to session. Once the experimenter presses “Confirm”, the session proper begins.

### 2.3 Settings files

Configuration settings files (**Figure 2B**) define numerous aspects of M-USE sessions (e.g. what experimental hardware is connected, or what tasks are available for participants to run) and task (e.g. how many trials are in a block, or termination criteria). They are loaded and parsed during the Session Level’s SetupSession state and each Task Level’s SetupTask state (**Figure 1B**) by the

ImportSettings module (**Figure 2A**). The session level files include an EventCodeConfig that allows customization of the codes that can be sent out by timing synchronization hardware such as our Arduino-based SyncBox (see Appendix), and a SessionConfig that governs other parameters, such as which tasks will be available to run this session. Up to six standard task level configuration files are automatically read if they exist, and developers can add other custom files if needed. These files enable numerous aspects of each task to be configured, including which variables will be presented on the Experimenter Display, enabling them to be modified in real-time. The M-USE Task Configuration File Reference (see Appendix) contains complete details for all Session, Task and Trial configuration files used in the Task Library.

### 2.4 Participant Input

Participant input is mediated through InputTracker modules (**Figure 2A**), for example MouseTracker, JoystickTracker, and GazeTracker modules handle input from computer mice, joysticks, and eyetrackers, respectively (**Figure 2D**). SelectionHandler modules (**Figure 2A**) integrate InputTracker values with details of the scene to determine if a selection has been made. The determinants of selection can vary widely: for instance, touch-based selections might require touching an object longer than a minimum and shorter than a maximum duration; gaze-based selection might only require gaze to be maintained on an object for longer than a minimum duration; while first-person joystick selection might require a participant to navigate through a scene and collide with an object to select it. Despite these very different inputs and selection conditions, any successful selection from a SelectionHandler is treated equivalently by M-USE, and in particular can be used to trigger the end of a trial state, making it easy for very different participant actions to have the same effects on the experimental state-flow logic.

### 2.7 Data files

M-USE’s data files are output into a hierarchical folder structure that allows the complete reconstruction of an experimental session (**Figure 2C**). Most of these are automatically generated by instances of the DataController class (**Figure 2A**), which allows developers to easily specify which variables are included, and when these variables’ current values need to be written to a data file.

The top-level Session data folder contains complete copies of the configuration settings folder (see 2.3) and Unity’s automatically-generated log file, a customizable overall session summary file, a simple timing document for the Session Level states, and a folder of data from the SelectTask state. Each run through SelectTask generates a new sub-folder, itself containing FrameData that primarily tracks frame-by-frame changes in participant input, as well as Serial Sent and Serial Received data files that record communication to and from the SyncBox, which can include event code and TTL pulse times for synchronizing with other hardware, commands to trigger reward pump delivery, and the output of photodiodes used for precise determination of monitor frame onsets. If eyetracking is active, SelectTask data will also include a GazeData file. Once a task begins, data is written to its subfolder, including single Task, Block and Trial data files, and additional folders for FrameData, Serial Sent/Received Data, and GazeData. These latter folders contain a file for each trial in the task, allowing for easy manual loading and visual inspection of data. Matlab processing scripts will parse an entire session’s data, creating single.mat files for each data type, for each task, for use in further analyses (https://github.com/Multitask-Unified-Suite-for-Expts/M-USE_Analysis).

### 2.8 Event Codes

Event codes enable post-hoc synchronization of M-USE data files with data files produced by other experimental devices (e.g. neural data acquisition and stimulation systems). Codes are handled using the EventCodeController, which communicates via the SyncBoxController and SerialPortController modules to communicate with the Arduino-based SyncBox (**Figure 2A, 2E**). Fully-customizable codes are specified in either Session or Task level EventCodeConfig files. Sent codes and their corresponding times are stored in the FrameData, SerialSent and SerialReceived Data files (**Figure 2D**), as well as in the files produced by whatever hardware receives these codes, enabling post-hoc timestream syncing. By default, codes are sent at the start of each state at the Session and Task level, as well as for stimulus appearance, fixation onsets to stimuli, selection of stimuli, registration of participant responses, etc.

### 2.9 Gaze Tracking

M-USE uses the EyetrackerController module (**Figure 2A**) to communicate with a Tobii Spectrum eyetracker (**Figure 2F**). When eyetracking is active (specified in the SessionConfig settings file), a Calibration state and corresponding Calibration child level is available at either the Session or Task level, which can be accessed as needed to perform initial calibration and later during a session to redo calibration or perform drift corrections. Activating an eyetracker also activates the corresponding GazeDataController, which stores its data in a series of trial-indexed files, similarly to the FrameData (**Figure 2D**).

### 2.5 Experimenter Display

The ExperimenterDisplay module (**Figure 2A**) controls an optional second display (**Figure 2G**) that provides ongoing information about participant performance and allows experimenters to make real-time adjustments to experimental variables. This display includes a panel that mirrors the participant display with task-specific overlaid information (gaze position if eyetracking is active, highlighting of correct choices, etc.); panels that give text summaries of performance over the most recent trial, block, task and session; a panel listing the available hotkeys; and a panel of variables that can be directly adjusted using sliders or the keyboard. Task developers have complete control over the information to be overlaid on the mirror view panel, the content of the text summaries, the specific hotkeys available, and the adjustable variables.

### 2.10 Selection and reward/token/slider feedback

A number of M_USE modules control customizable feedback to the participant in the form of selection feedback or reward/token feedback (not shown in Figure 2). Three types of selection feedback can be presented upon object selection: auditory (the specific sound and duration are customizable as needed), a translucent halo displayed above and around a selected object (with customizable colour, size, and duration), or a 2d image displayed at the selected location (customizable image and duration). Control over *selection* feedback is deliberately separated from *reward* feedback modules, which include slider and token style visual feedback combined with customizable audio feedback, allowing experimenters to design tasks where rewards are temporally separated from immediate choices (Boroujeni et al., 2022).

### 2.11 Frame Detection

To enable complete reconstruction of an experimental session with sub-ms timing precision, the user can record data from light sensors positioned above two small patches on the participant monitor (**Figure 2C**) that are connected to the SyncBox (**Figure 2E**). An example of light sensors mounted onto 3D-printed clamps is provided (see Appendix). These panels are controlled by the FlashPanelController module (**Figure 2A**), such that one “timekeeper” panel flips every frame, while the other “reconstruction” panel goes through a unique 24-frame cycle (treating black frames as 0 and white frames as 1, it counts from 0 to 7 in binary in 3-frame ‘digits’, resulting in the complete sequence 000 001 010 011 100 101 110 111). The timekeeper panel is used to reliably find the physical onset time of each frame, providing an accurate timing that goes beyond Unity’s intrinsically reported frame onsets. Additionally, the reconstruction panel allows identifying “stuck” or “skipped” frames. The Experimenter Manual and accompanying analysis scripts describe in detail how the timing analysis automatically corrects the frame onset timing of FrameData files (see Appendix, and section 2.12).

### 2.12 M-USE’s temporal precision, synchronization of data streams

Temporal precision to the level of the nearest frame (16.7 ms on a standard 60 Hz commercial monitor) is obtainable by default in M-USE, both for real-time control and for post-hoc data reconstruction. The entirety of M-USE’s state logic inherits from Unity’s frame-locked Update(), FixedUpdate(), and LateUpdate() classes, whose *order* within a frame is reliable but whose precise *timings* within that frame are effectively uncontrollable. This means that experimenters can be confident that any desired methods will be run during a frame, provided the frame is displayed at all.

Each data file associated with a control level (Session, Task, and Trial data files) contains the frame numbers of the initialization and termination of each state in that level, which correspond to the same frame numbers provided in the FrameData files. FrameData includes the event codes sent on each frame, which can then be used to synchronize any data from equipment that receives these event codes to the frame level. Syncing to a sub-ms level requires further processing. The Synchbox samples from the light sensors above the monitor light sensors at 300 Hz. The signal for each photodiode is filtered using a zero-phase Butterworth bandpass filter with frequency cutoffs of 1Hz to 75Hz. The filtered signal is mean-normalized using a sliding window every second and interpolated to a 1500 Hz sampling rate using spline interpolation for the left photodiode and a shape-preserving piecewise cubic interpolation for the right photodiode. In a next step the transition points from the left photodiode are identified by detecting peaks and troughs of the signal that are separated by 8 ms (shown in red and blue respectively in **Figure 3A**). The right photodiode signal was binarized first, and then transition points were identified by finding falling and rising points using the first derivative of the signal (shown in red and blue respectively).

**Figure 3.**
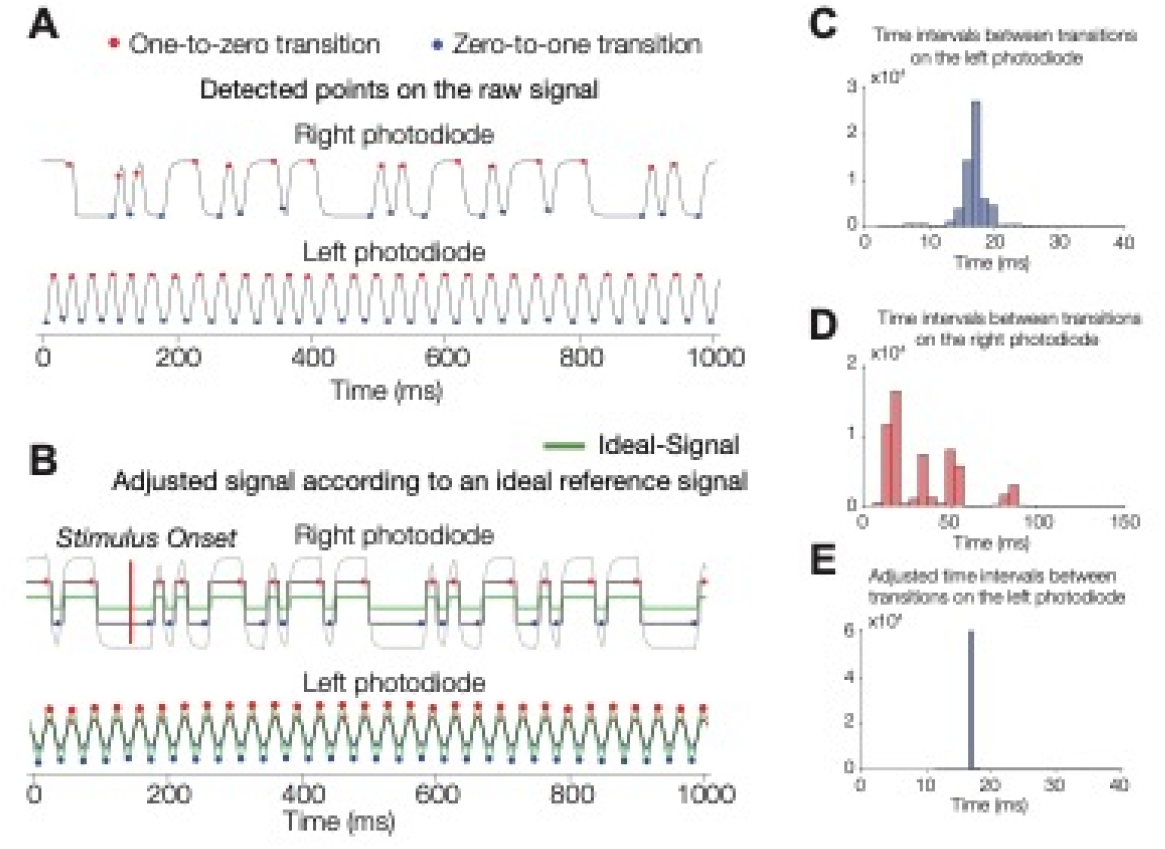
Temporally precise reconstruction of frame onsets. To precisely reconstruct events to the physical monitor frame at which events they occurred during task performance, photodiodes can be used to track white-black flashing sequences. Analysis scripts are provided on github to detect the measured onsets (A) and adjust the onset times in case frames were stuck of delayed (B). The time intervals between flashes on the right diode (C) and left (D) diode are precisely adjusted (E).

We then used a sliding window that reconstructed ideal signals and cross-correlated them with the recorded signals from the left and right photodiodes. The correlation coefficient value and sample lag for the best fit in each sliding window were returned. For all sliding windows that cover the entire right photodiode signal, we used the first derivative of the lags to detect sections with potential skipped or stuck frames. To determine the exact frame discrepancy, the continuation of the previous window without any shift was subtracted from the actual signal and the first non-zero location that lasted at least one frame was determined. Through this analysis, the full-frame transitions on the monitor can be reconstructed and compared with the unity frame data. Finally, to time-rectify the event codes, we extracted the recorded event code time from the Arduino and identified the nearest preceding frame transition from the left photodiode.

### 2.13 Server-based configuration reading and data writing

M-USE includes a ServerManager module that enables access to remote servers via HTTP requests, and the server-side php scripts used to handle these requests. This is integrated with both the SettingsManagement and DataController modules, allowing reading of configuration files from, and writing of data files to, such servers. The server address, the option to use a server-based settings folder, and the option to write data to the server, are specifiable by the experimenter in the Initialization Screen.

### 2.14 Web GL Builds

M-USE can produce Web GL builds (see http://m-use.psy.vanderbilt.edu/play/) that can be accessed using any standard internet browser from most modern computers and tablets (but not phones). For the most part, these builds perform identically to native Windows or Mac builds, with one critical difference being that they cannot access local hard drives. This difference means that session files are either read from a remote server or from the defaults stored in the build itself, and data files can only be stored on a remote server.

### 2.15 Common Multi-dimensional Objects

Multiple tasks in M-USE use multidimensional objects, so called Quaddles, initially described in (Watson et al., 2019a), and extended for the M-USE software suite (see Appendix). These vary in more than 10 different feature dimensions (body shapes, arm shapes and tilt, colors, patterning of the body, etc.), each of which can have many different values (specific shapes, colors, etc). Quaddles are automatically generated with the freely-available software Blender.

### 2.16 Task Paradigms Implemented in M-USE

M-USE’s Task Library contains seven pre-configured tasks with unique task and trial states (**Figure S3**): Visual search, flexible learning / attentional set shifting, visuospatial problem solving, spatio-temporal (‘what-when-where’) relational memory (implemented using an object sequence learning task), effort control and motivation, working memory, and a continued recognition (also called visual working memory span). Video demonstrations of these tasks are available online (http://m-use.psy.vanderbilt.edu/tasks-3/). They were chosen to assess higher cognitive and motivational functions that depend on different anatomical subfields of the prefrontal cortex and their associated network connections to the medial temporal cortex and hippocampus, the amygdala, the basal ganglia, and the parietal and temporal cortex **Figure 4A,B** (Passingham, 2021). Performance of these tasks allows evaluating well-defined behavioral metrics that are relevant for assessing cognitive constructs of attention, working memory, cognitive flexibility, motivation, and relational long-term memory (**Figure 5**).

**Figure 4.**
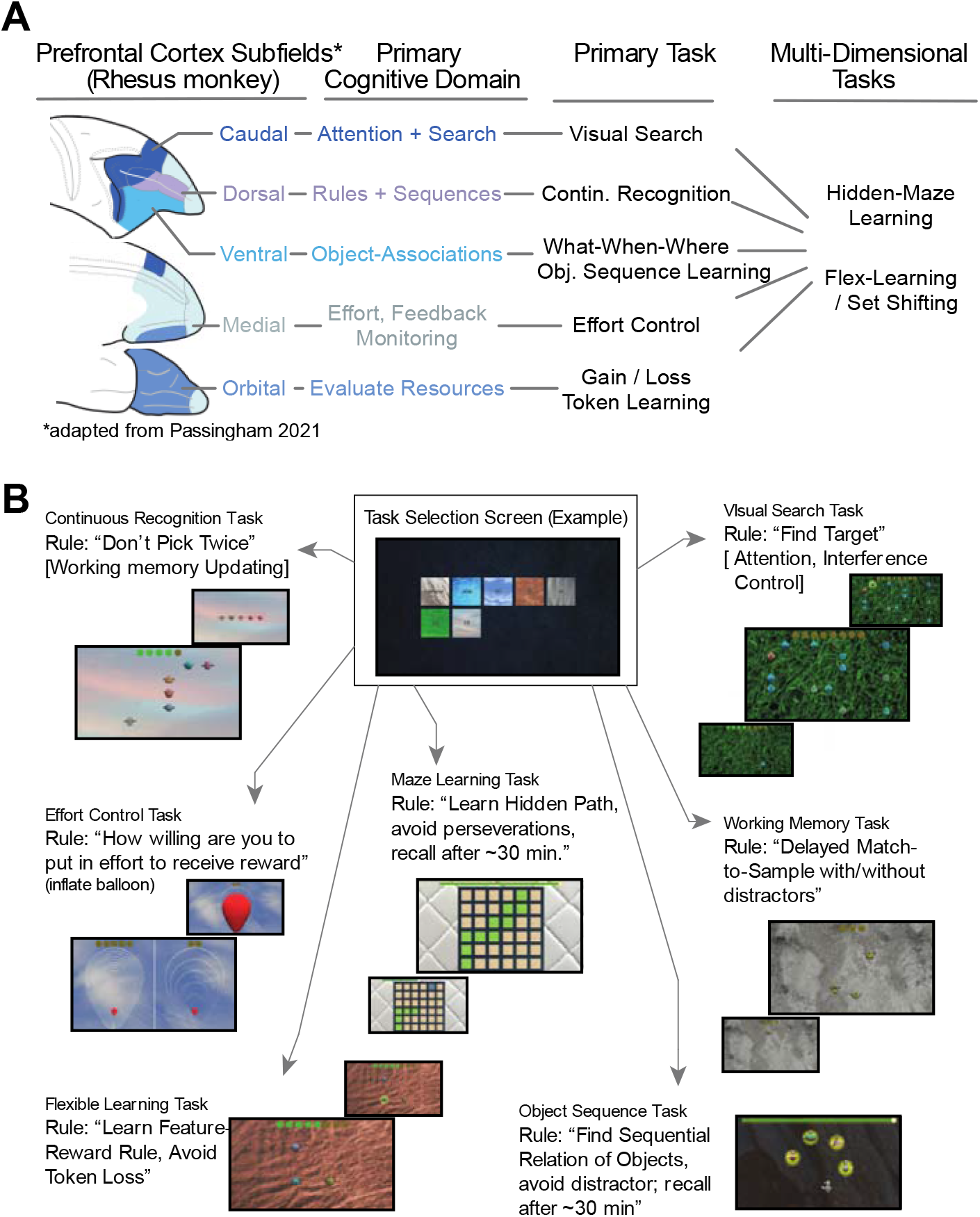
Overview of tasks (A) M-USE state systems incorporates tasks that were chosen to assess functional domains realized by separate subfield of the prefrontal cortex according to Passingham (2021) who distinguished five cortical subfields (left) and primary functional domains. We measure these domains with five primary tasks and two multidimensional tasks (right). (B) Depiction of the multi-task selection screen subjects use to choose tasks (top middle), and illustrations of the tasks and task rules.

**Figure 5.**
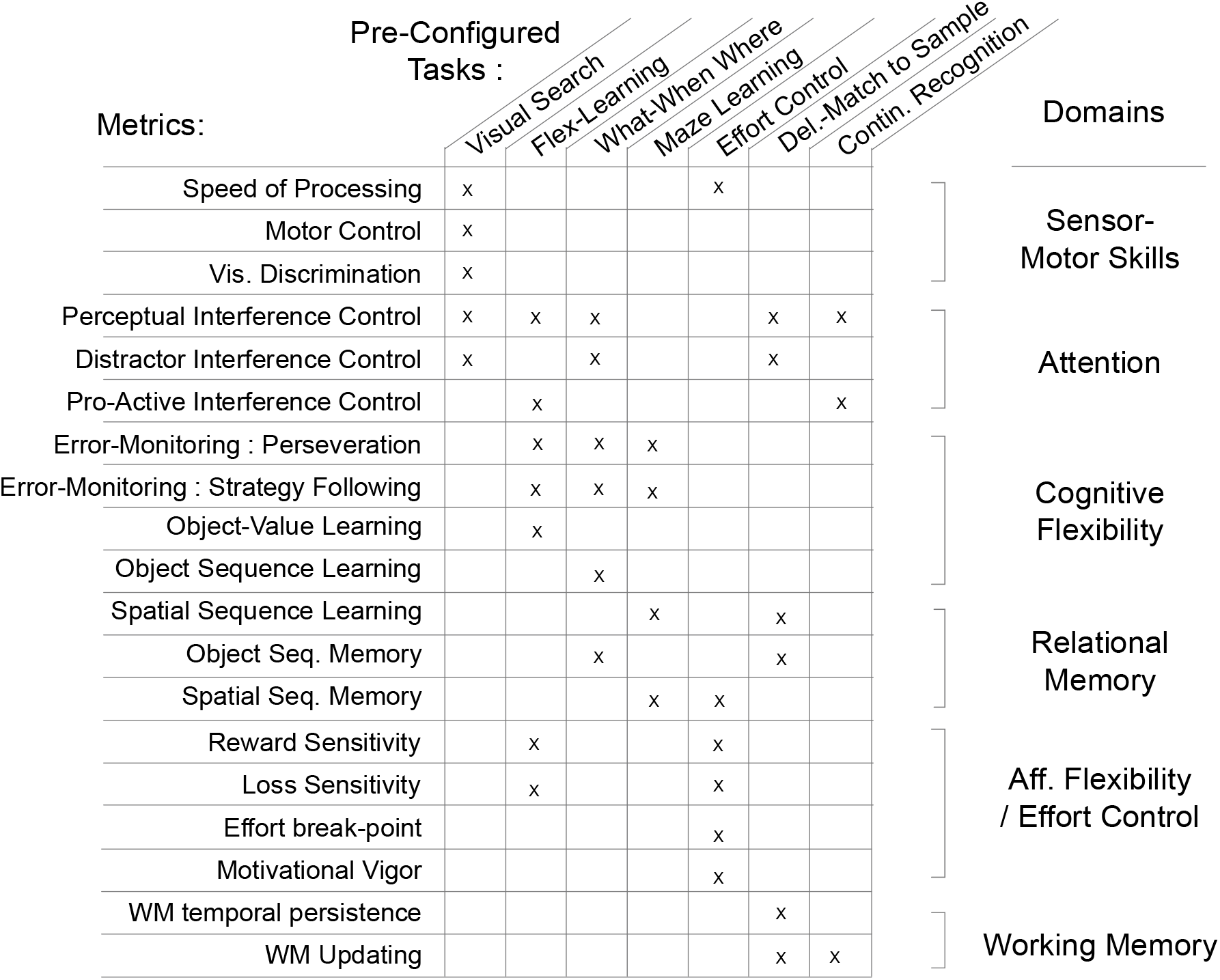
Performance Metrics of pre-configured M-USE tasks. The pre-configured tasks of M-USE (columns) measure a wide array of performance metrics (rows). The metrics map onto six different cognitive affective domains that encompass RDoC constructs (rightmost column). How the metrics are calculated is described in the main text and realized in matlab scripts (The MathWorks, Inc.) accompanying the M-USE github (https://github.com/Multitask-Universal-Suite-for-Expts).

### 2.16.1 Visual Search

### Purpose

The Visual Search (VS) task evaluates focused attention, control of distractibility (how much detection is slowed with increasing distractor number and similarity), and speed of processing (baseline target detection time).

### Description

VS measures how fast and accurate subjects are in detecting a target object among distractors. The task varies the number of distractor Quaddles and their perceptual similarity to target Quaddles (defined as the number of shared feature values). Our VS task is run in one block of 200 trials. In the first ten trials, a single target object is shown at random locations and the subjects is rewarded for choosing it. In the subsequent 190 trials, the target object is shown together with 3, 6, 9, or 12 other multidimensional distractor objects, which share 1, 2 or 3 features with the target, determining their similarity.

### Relevance

VS task in M-USE are standardly used to measure attention and search processes (Wolfe et al., 1989; Wolfe, 1992), which are compromised to different degrees in major neuropsychiatric disorders (Millan et al., 2012). These processes are behaviorally dissociated from working memory capacity (e.g. Meier & Kane, 2013) and closely associated with the lateral prefrontal cortex and the dorsal fronto-parietal attention network (Passingham, 2021). Rhesus monkeys with disruptions of the lateral PFC, the frontal eye fields, or intraparietal cortex are impaired at covert selection of a visual target stimulus among distractors (Keller et al., 2008; Rossi et al., 2007; Wardak et al., 2006). Visual search performance is modulated by cholinergic neurotransmission with cholinergic-esterase inhibitors enhancing distractor filtering (Hassani et al., 2021). In addition to selective attentional processing, overall accuracy of searching for a target requires working memory (Poole & Kane, 2009), and indexes an overall ‘speed of processing’ which is a metric that is consistently correlated with deteriorating executive control functioning and fluid intelligence of the aging brain (Ball et al., 2007; Edwards et al., 2002). Changes in both, distractor filtering and speed of processing, are prominent signatures of cognitive aging (Kennedy & Mather, 2019).

### Metrics

We compute four VS metrics (**Figure 5**). Accuracy and search reaction times are quantified for each distractor condition and reduction of accuracy and slowing of target detection with increasing numbers of distractors is fit with a linear regression model. The regression intercept indexes *speed of processing* (1), while the slope of the curve reflects how strong accuracy decreases with additional distractors with a shallower slope indexing better attentional filtering of distractions, i.e. *distractor interference control* (2). The difference between search slopes for displays with high versus low target-distractor similarity indexes *perceptual interference control* (3), while the difference between their intercepts indexes an overall skill of ‘visual discrimination’ (4).

### 2.16.2 Flexible Learning / Attentional Set Shifting

#### Purpose

The Flexible Learning (FL) task assesses how fast and accurate subjects learn new attention sets and how sensitive subjects are to gains and losses. It measures various cognitive constructs underlying cognitive flexibility including the degree of perseverative responding, and the costs associated with switching attention sets from one rewarded feature to a new feature of the same feature dimension (intra-dimensional shifts) or of a different feature dimension (extra-dimensional shift).

#### Description

On each trial, three Quaddles are presented at random locations equidistant to the screen center, along with a token progress bar with eight slots for tokens at the top of the screen. Subjects have to identify through trial-and-error which of three objects lead to most tokens. In blocks of ∼30 trials token gain is associated with one visual feature found on only one object (e.g. the colour red), while the other two objects have other values along this feature dimension (e.g. different colors), and all three objects vary in features of another feature dimension (e.g. different body shapes). Thus, on every trial there is one target feature and five irrevelant features. Within a block the set of 3 objects stays constant.

Blocks differ in four ways. The object set can either be the same as in the previous block, or different (*Same* vs *New* block switch), and the rewarded feature can either be from the same feature dimension as the previous block or from a different feature dimension (*intra-dimensional/ID* vs *extra-dimensional/ED* block switch). Blocks also differ in reward schedules: participants gain either two or three tokens for correct choices (G2 vs G3), and lose one or three tokens for incorrect choices (L1 vs L3).

Blocks always start with a token progress bar that contains three tokens, enabling participants to determine after the first error whether it is a L1 or L3 block, and after the first correct choice whether it is a G2 or G3 block. The token progress bar contains 8 slots that need to be filled with tokens before subjects ‘cash out’, e.g. receive juice reward. The task quantifies the number of trials needed to reach a learning criterion (trials-to-criterion) and it measures different error types that indicate whether subjects persevere on previous reward rules or follow suboptimal strategies (proportion of errors that follow errors). The FL task proceeds through a user specified number of blocks.

#### Relevance

FL combines aspects of four types of tasks used in the literature to assess functions of the lateral prefrontal and orbitofrontal cortex and the basal ganglia. *First*, it requires learning the value of features, similar to learning feature-based rules in the Wisconsin Card Sorting Task or the Category Set Shifting Task (Moore et al., 2013). Learning such rules requires choosing objects on the basis of their features, and determining whether to continue choosing the same feature after feedback (e.g Win-Stay, Lose-Switch), functions causally supported by lateral and ventral prefrontal cortex and considered a marker of age-related cognitive changes (Buckley et al., 2009; Jang et al., 2015; Rudebeck et al., 2017). *Second*, the intra-vs extra-dimensional shifts resemble classical ED/ID attentional set shifting tasks that quantify whether subjects have higher switch-costs (learn slower) when a new target is from a different target feature dimension (Brown & Tait, 2016; Roberts et al., 1988). Higher switch costs at ED than ID indicate that top-down control is hierarchically structured, because it will be easier to switch the target representation at the same ‘intra-dimensional’ level of a hierarchy. *Third*, the FL task measures the degree of pro-active interference from the memory of objects in previous trials. Learning will be slower when a feature found on distractors in the previous block becomes the target feature in a new block (reflecting latent inhibition) compared to target features that are novel. The degree of latent inhibition will be less in subjects with stronger cognitive control. *Fourth*, and finally, the FL task evaluates whether higher incentives (G3 vs G2 blocks) lead to faster learning, which indicates sensitivity to positive valanced reward. Sensitivity to positive outcomes is diminished in subjects diagnosed with major depression (Vrieze et al., 2013). Similarly, the FL task evaluates whether higher losses lead to decreased learning, which indexes how sensitive a subject is to aversive loss outcomes is enhanced in subjects diagnosed with anxiety.

#### Metrics

Performance of the FL task provides five metrics (**Figure 5**). The speed of ‘feature-value learning’ (1) corresponds to the time to reach criterion performance defined as the first trial leading on average to 75% correct performance over a forward looking 10-trial window. Learning speed indexes cognitive flexibility and comprises multiple subfunctions that this and other tasks tap into. Comparison of the learning speed for blocks with different gains indexes reward sensitivity (2) and for different losses indexes loss sensitivity (3). Flex-Learning allows quantifying error monitoring for avoiding perseveration when choosing repeatedly non-rewarded object features (4). A previous-trial analysis that measures the accuracy in the trials after an error trial (EC_n_ analysis) quantifies a perseveration score that is higher when a subject shows a shallow rise or no improvement of performance after errors. Comparing of the perseveration score in blocks with the same versus novel objects than in the previous block estimates the vulnerability to ‘pro-active interference’ from the previous blocks target object (5).

### 2.16.3 Object Sequence Learning Task

#### Purpose

The Object Sequence learning task (also denoted *What-When-Where/WWW* task) is an extension of an item-item paired associates task that measures relational memory for objects and contexts, error monitoring, the use of efficient learning strategies, and interference control of filtering distraction.

#### Description

The Object Sequence learning task shows the subject six objects and a progress bar on top of the screen. Five of the objects are part of a temporal object sequence the subject has to learn by touching individual objects in a prespecified order. The sixth is a distractor that is not part of the sequence, but shares several features with the object in either position 2 or 4. A colored halo around the touched object provides touch and accuracy feedback (yellow for correct, gray for incorrect), as does a tone (high pitched for correct, low for incorrect). When subjects complete a sequence, the slider will have advanced through the entire extend of the progress bar which then flashes and either fluid reward is dispensed (for nonhuman primates) or the task score is incremented (for human subjects). Different object sequences are shown on unique context background images. Each sequence is repeated at a later time point within a session with the same objects and identical context background. Faster learning of the repeated than initial session quantifies relational memory of object sequences.

#### Relevance

The Object Sequence Learning task resembles classical paradigms assessing in nonhuman primates the learning of sequential relationships of photographs (Harlow, 1949) and objects from different categories in the ‘simultaneous chain task’ (Altschul et al., 2017; Terrace, 2005). Learning and inferring the order of objects depends in primates on the ventrolateral prefrontal cortex (Petrides, 2005) and the hippocampus (Heuer & Bachevalier, 2013), which has been considered essential for learning non-spatial and spatial sequences (Buzsaki & Tingley, 2018).

#### Metrics

Performance of the object sequence learning task provides four metrics (**Figure 5**). The average number of trials required to complete a sequence quantifies object sequence learning abilities (1), which involves forming object-object associations and chunking them into a sequence. Errors differ depending on whether an object from earlier positions is erroneously chosen again (*repetition errors*), or whether subjects chose the wrong object at a particular temporal position (*slot errors*). The proportion of repetition errors to all errors signifies the ability of ‘error monitoring/perseveration’ (2), while the proportion of non-repetition slot errors indexes error monitoring/strategy following (3), and the proportion of retouch errors (erroneously not retouching the last correct tile after an error) indexes difficulty maintaining the task rule. Object sequences are accompanied by one distractor object that shares features with one of the objects. The number of distractor errors at the ordinal position at which the distractor was similar to the correct object is divided by all distractor errors to quantify how likely the distractor is confused with the correct object that specific to the ordinal position, i.e. it measures ‘temporal distractor interference’ (4). This ordinal positioning distractor effect is similar to the symbolic distance effect used in the literature to infer the learning of sequential relationships (e.g. Orlov et al., 2000).

### 2.16.4 Maze Learning Task

#### Purpose

The Maze-Game Learning task evaluates how fast and accurately subjects learn hidden trajectories through a grid-based maze. This requires visuo-spatial learning and memory skills as well as the use of task rules and feedback (error monitoring).

#### Description

The Maze Learning task requires subjects to find a hidden maze’s trajectory through a 6×6 grid that is uniform except for highlighted start and end points. Trajectories are from 10-15 tiles long, and contain from 3 to 6 turns. Subjects initiate the task by touching the starting tile, and make choices by touching tiles horizontal or vertical relative to their last tile. After errors, subjects must re-confirm the last correct tile by choosing it again. For every correct choice the progress bar at the top of the screen is advanced. The slider completes the progress bar when a choice connects the last tile of the trajectory to the colored end point. Each maze has a unique textured background so that some of the mazes that are learned early in a session can be repeated at a later time in the same session to facilitate testing longer term retention of the initially learned spatial path.

Correct choices are signaled by a brief (duration *****) green flash of the chosen square and a high-pitched tone. Errors are signaled with a low-pitched tone and other tile colors: black for a “rule-abiding” horizontal or vertical choice that is not on the maze, red for a “rule-breaking” choice of a non-adjacent tile, green/grey flashing of the last correct tile if subjects repeatedly fail to press it after another error.

#### Relevance

The Maze task is a variation of the Groton-Maze-Learning task (GMLT) paradigm used in humans to measure rule-based and spatial learning (Pietrzak et al., 2008), and is considered to have particular high cross-species validity for understanding the formation of spatial maps (Redish et al., 2022). Performance scores of the maze task are correlated with problem solving ability and fluid intelligence. The task has high demands on error monitoring, learning from errors, and spatial working memory which indexes problem solving skill. Humans with schizophrenia show impairments of spatial learning (Bakker et al., 2020), error monitoring (number of rule-breaking errors), efficiency (time and number of moves to completion), and show perseverations (Snyder et al., 2008) which can improve with dopaminergic and cholinergic drugs (Lieberman et al., 2013; Pietrzak et al., 2009; Pietrzak et al., 2010). Maze learning also involves the recall of spatial sequences, which involves the hippocampus (Buzsaki & Tingley, 2018), ventrolateral prefrontal cortex (Axelsson et al., 2021) and the dorsolateral prefrontal cortex (Mushiake et al., 2006; Saito et al., 2005). It shares similarities to the path learning of nonhuman primates controlling cursors (Mushiake et al., 2001), and spatial relational learning of rodents in the Morris Water Maze, which is linked to the hippocampal system and to cholinergic modulation as muscarinic compounds reverse deficits in spatial learning and memory (Popiolek et al., 2019).

#### Metrics

Performance of the Maze-Learning task provides four metrics (**Figure 5**). The average number of choices to complete a maze, normalized by the path length of the maze quantifies the speed of ‘spatial learning’ (1). Error monitoring is indexed by the proportion of rule-breaking versus rule-abiding errors (indexing strategy maintenance) (2), and by the proportion of repetition errors (indexing perseveration) (3). Performance on mazes that are repeated with their unique context background and start-/end-position within a session evaluate spatial memory (4).

### 2.16.5 Effort Control and Motivation

#### Purpose

The Effort Control (EC) task measures how motivated subjects are to work for a reward. It indexes sensitivity to costs (effort) and to reward value (benefit), and how cost-benefit ratios motivate behavior.

#### Description

The EC task begins with two sets of balloon outlines, one on each side of the screen, and participants must choose one set. The number of outlines can differ between sides, and indicates the workload required by the participant to inflate and pop the balloon and receive their reward. Thus the number of outlines corresponds to the effort or costs of each set. A number of coin-like tokens are presented just above each balloon choice, which corresponds to the amount of reward that will be given after the balloon is inflated. Costs and reward ratios are systematically varied in order to evaluate their motivational effects.

#### Relevance

Effort control deficits are a hallmark of multiple psychiatric disorders (Der-Avakian et al., 2016) and higher levels of motivation are major predictors of outcome success for treatments in patients with schizophrenia and depression (Berwian et al., 2020; Gold et al., 2013). The EC task measures motivation as a costs / benefits ratio threshold, and the threshold which effort is withheld reflects the motivational *’break point*’ in operant tasks using progressive ratio schedules (Hodos 1961, Roane et al., 2001). The EC task differs from classical progressive ratio tasks by allowing subjects to choose between options with higher-effort/high reward versus options with lower-effort/lower-rewards, and by indexing effort as the number of touches required to reach a goal as opposed to the numbers of lever presses or climbing taller barriers for reaching rewards (Salamone et al., 2007; Salamone et al., 1997; Salamone et al., 2009). Effort control is causally supported by activity in the dorsal anterior cingulate cortex in humans and nonhuman primates (Passingham, 2021), is associated with striatal dopamine synthesis (Westbrook et al., 2020), can be strengthened using amphetamines in individuals with low motivation (Cocker et al., 2012), and is predicted by higher activity of noradrenergic neurons in the locus coeruleus (Bornert & Bouret, 2021).

##### Metrics

Performance of the Effort-Control task provides three metrics (Figure 5). The motivational “break point” where subjects reliably choose the lower effort option (1) can be expressed in terms of effort difference between the two choices (difference in the number of outlines on each side) or simply in terms of the effort associated with the harder choice. The break-point computation also entails an estimate of (2) the subjective value of the chosen side as an ‘expected utility’ metric that reflect the ratio of expected benefits (amount of tokens) and costs (number of balloon outlines) (Amemori & Graybiel, 2012; Levy et al., 2010). Finally, the average time taken to inflate a balloon normalized by the number of touches needed provides an estimate of “motivational vigor” (3).

#### 2.16.6 Delayed-Match-To-Sample Working Memory

##### Purpose

The Working Memory (WM);4task tests how well subjects sustain visual objects in working memory over delay. The task varies the delay duration to measure the temporal fidelity of working memory, the complexity and similarity of objects to measure the degree of perceptual interference, and the presence of distractors to measure memory interference.

##### Description

The WM task is a classical delayed-match-to-sample task. A trial begins with a long presentation of a single sample object at a random location for 0.5s. After a delay, two or three test objects are shown at random locations, and the subject should touch the object that matches the sample. The task has variable delay durations (e.g. 0.7, 1.5 or 3 seconds), 0 or 3 distractors shown during the delay period, and test objects that share variable numbers of features with the sample.

##### Relevance

Working memory abilities mediate performance of many other tasks (e.g. feature learning) and thus are connected to performance variations across multiple tasks (Collins et al., 2014; Collins & Frank, 2013; Unsworth & Robison, 2017).

##### Metrics

WM results produce two metrics: choice accuracy as a function of delay time quantifies the temporal persistence of visual memory, and choice accuracy as a function of distractor presence quantifies its susceptibility to distractors (**Figure 5**).

#### 2.16.7 Continuous Recognition and Self-ordered Working Memory

##### Purpose

The Continuous Recognition (CR) task is a variation of visual working memory span tasks and measures how well subjects dynamically update and maintain in working memory of increasing numbers of objects.

##### Description

The CR task displays multidimensional objects on the screen and requires subjects to choose an object they haven’t previously chosen. On successive trials the total number of objects increases such that the display contains novel (N), previously-chosen (PC), and previously-viewed but not-previously-chosen (NPC) objects. When subjects correctly choose an N or NPC object, they receive a visual token that fills a token bar on the top of the screen. When the subject choses an object that was already chosen in any of the previous trials, the block ends and feedback is provided by showing all previously chosen objects in the order they were chosen, highlighting the last, wrongly chosen object together with the first instance it was presented in a red colored frame.

##### Relevance

The CR task requires the continuous monitoring and updating of working memory contents, two key functions underlying the ‘cognitive flexibility’ construct (Uddin, 2021). The updating of working memory content is well separated from processes linked to response inhibition (Chase et al., 2008; Friedman et al., 2006; Miyake et al., 2000), and is causally supported by the ventral and dorsal lateral prefrontal cortex in humans and NHPs (Petrides, 1991; Wager & Smith, 2003). The CR task has the same requirements as the Delayed Recognition Span task that depends on prefrontal cortex and its dopaminergic and noradrenergic functioning (Moore et al., 2005). It requires self-ordered search for a new target, which has been associated specifically with the ventral and lateral prefrontal cortex, but not the orbitofrontal or parietal cortex (Axelsson et al., 2021; Champod & Petrides, 2007; Walker et al., 2009), and in humans is causally supported by the inferior frontal gyrus (Chase et al., 2008). Deficits in self-ordered selection tasks are evident in humans diagnosed with attention-deficit-hyperactivity disorder (ADHD) (Dowson et al., 2004; Fried et al., 2015) and schizophrenia (Badcock et al., 2005; Pantelis et al., 1999), including first degree relatives and monozygotic twins of schizophrenic patients (Pirkola et al., 2005; Wood et al., 2003) even when they have no altered cognitive functioning (Joyce et al., 2005). Pharmacologically, self-order recognition memory depends on the acetylcholinergic subreceptors of the muscarinic type (Callahan, 1999; Lange et al., 2015; O’Neill et al., 2003; Schwarz et al., 1999).

##### Metrics

Performance of the Continuous Recognition task measures the ‘WM updating’ ability indexed as the number of correctly chosen objects before committing an error. The updating metric can be extended by computing a novelty updating bias inferred from the difference of the proportion of correctly chosen novel (N) versus not-previously chosen (NPC).

#### 2.16.8 Touch Hold Release (THR) task

##### Purpose

The Touch-Hold-Release task (THR) teaches subjects the fundamentals of using a touchscreen with M-USE tasks. Subjects learn to touch, maintain touch (hold) and release objects on the screen, an must maintain holds for longer than a minimum and shorter than a maximum duration in order to receive reward. Thus THR trains consistent and reliable touch behavior, particularly important for NHPs.

##### Description

The task displays a square on the screen that blinks from white to blue. Touching the blue square for the correct duration results in positive feedback. Touching the white square, or touching the blue square for the incorrect duration, will result in negative feedback. The difficulty level of the THR increases by reducing the square size and randomize its positioning on the screen.

##### Relevance

The THR task is primarily meant as a training exercise, whose key performance metric is accuracy. When an acceptable level of performance is maintained, NHP subjects are ready to begin training in the other tasks. It could be extended to provide insights into motor control when accuracy is evaluated for differently sized squares.

## 3. Results

### M-USE is a versatile multitask programming platform

M-USE establishes a platform for controlling multiple cognitive and motivational tasks with a modular architecture (**Figure 1A**), a hierarchical state logic for organizing information flow at session, task, and trial levels (**Figure 1B**), and a versatile organization of multiple inputs and outputs (**Figure 2**). These platform features facilitate customizing, extending, and developing new tasks and features into M-USE. M-USE is *multi-task*, enabling multiple tasks to be run within single sessions, and *multi-build*, enabling similar studies to be run in different setups, including on web-servers. Its stimulus handling modules can flexibly present 2D or 3D rendered stimuli. Its selection (choice) handling module treats different physical inputs equivalently, enabling object selections by gaze, touch, button, and joystick, amongst others. The experimenter display provides real-time information on subjects’ performance, and allows key variables to be adjusted on a trial-by-trial basis. The modular programming of these features allows a developer with intermediate C# experience to program and establish a new task from scratch within a few days. For example, one of the authors (NT) required approximately fifteen hours to develop a classical cognitive control task to be functional and integrated into the multitask structure of M-USE. A knowledgeable programmer without prior Unity and C# experience who aims to code a M-USE task will need to first acquire Unity’s basic functionality, which undergraduate students have accomplished in ∼2-4 weeks, followed by another ∼2-4 weeks for reviewing the M-USE Developer’s Manual (*see* Appendix).

### M-USE allows assessing multiple cognitive constructs in single experimental sessions

The M-USE platform has been designed to evaluate rhesus monkeys and humans across multiple cognitive and motivational constructs in single behavioral testing sessions. M-USE accomplishes this using seven pre-configured tasks that probe cognitive and motivational functions, which causally rely on segregated prefrontal cortical subfields as established in lesion studies (Passingham, 2021) (**Figure 4**). The pre-configured tasks provide multiple behavioral metrics that evaluate (1) sensorimotor-skills (speed of processing, motor control precision), (2) attention (interference control), (3) cognitive flexibility (set shifting and error monitoring, which includes perseveration and strategy following), (4) memory (object-object sequence memory, spatial maze memory), (5) affective flexibility and motivation (effort control), (6) valuation of gains and losses; and (7) working memory (persistence and updating of short-term memory content) (**Figure 5**). We document the functionality and feasibility for assessing these domains by analyzing performance of nonhuman primates (NHPs, rhesus monkeys) trained on the preconfigured tasks. The NHPs performed up to five tasks each day, in sessions spanning 90-150 minutes. We report the results from two monkeys who had experience with the Visual Search task and the Flex-Learning / Set Shifting tasks prior to being trained on the spatial Maze Learning task, the Effort Control task, the Object Sequence Learning task, and the Delayed Match-to-Sample (Working Memory) task. The animals were gradually exposed to these tasks such that by the end of training, individual sessions contained up to five tasks. The order of the tasks was initially fixed with the newer tasks performed earlier in a session to ensure subjects learn these tasks prior to performing already trained tasks.

We found that sessions with 200 Visual Search trials were sufficient to measure the classical pop-out and set size effects: no effect of distractors on search times when target-distractor similarity is low (so called pop-out effect), but a gradual slowing with increasing numbers of distractors on high similarity trials (so called set-size effect, **Figure 6A**). These effects index the speed of attentional processing and efficiency of attentional control over distractor interference. Sessions of 150-180 trials were enough to index the temporal maintenance of a target object in working memory, evident in above-chance performance for delays up to 1.25 s and a decline to near chance performance at 1.75 s (**Figure 6B**). The working memory task additionally measures the representational fidelity of short-term memory content by introducing distractors in the test display that shared or did not share features with the probe (target) stimulus. We found a systematic decline in performance when the test display has high similarity with the target probe (**Figure 6B**). We introduced new maze paths with variable length and number of turns (path complexity) for the Maze Learning task in each session, which challenged the monkeys to learn the spatial trajectory and to avoid rule-breaking errors. We found that monkeys gradually decrease overall and perseverative errors when repeating a maze path on successive trials, documenting successful error monitoring, rule following, and spatial learning (**Figure 6C**). The Flex-learning / Set Shifting task requires learning feature-reward association in blocks of 25-40 trials with un-cued intra-and extra-dimensional (ID / ED) block switches. We found that learning speed (average number of trials to reach performance criterion), and ED and ID switch-costs are measurable with sixteen and more block transitions per session, requiring approximately twenty to thirty minutes of testing (**Figure 6D**). We found that monkeys learn the Effort Control task within one week. Running 80 trials in a session proved enough to estimate how likely the subject choosed the higher rewarded option (reward sensitivity) and how likely they prefer options with lower effort requirements (i.e. with less balloon outlines), indexing the subjects effort control level (**Figure 6E**). For training and evaluating object-sequence learning, we adopted a protocol that shows the same set of four objects, together with a fifth distractor objects, for ≥40 trials each session. We find that monkeys learn the 4-object sequence by gradually reducing erroneous choices of not yet learned objects (exploratory errors), but more prominently by learning not to touch objects from earlier in the sequence (retrospective errors), which indicates they gradually overcome difficulties to remember the learned sequence in working memory (**Figure 6F**). When the distractor is similar to the correct object at a particular time slot, subjects are more likely to erroneously choose the distractor in that slot, which is particularly apparent when a learned object sequence is repeated (**Figure 6F**). Successfully ignoring a distractor indicates control over interference, and thus error rates can index this ability.

**Figure 6.**
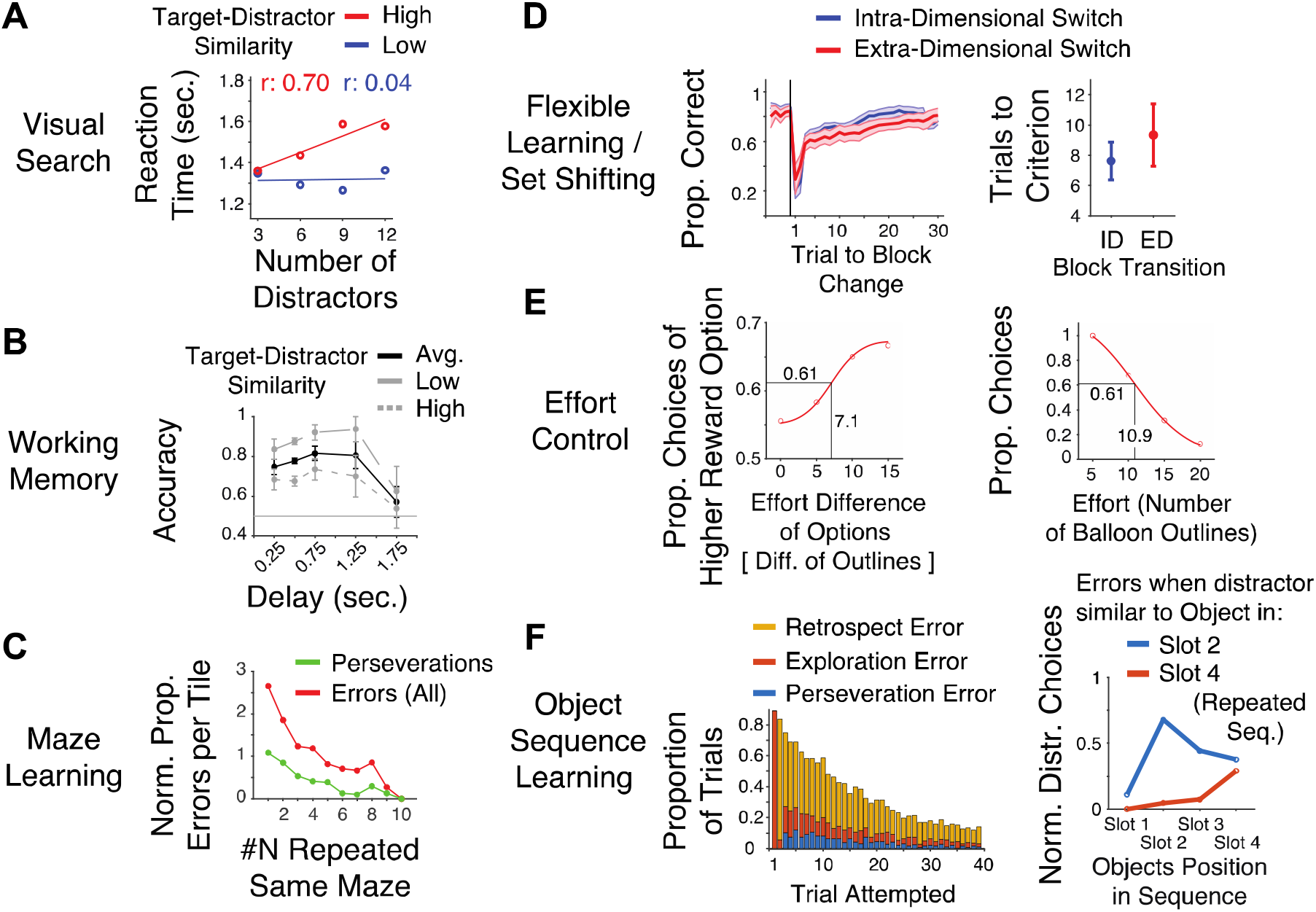
Extracting performance metrics from monkeys performing ≥3 of the pre-configured tasks per daily session. (A) Visual search regression slopes from 200 trials of a single session showing increased distractor interference (i.e. a conjunction search effect) evident in longer reaction times with more distractors (x-axis) sharing perceptual similarity with the target (red line: r=0.7 regression slope). Distractors that are dissimilar to the target cause a pop-out effect (blue line). (B) Working Memory: Delayed-match-to-sample performance across seven sessions using 150 trials each shows a drop in accuracy at 1.75 s delay and purer performance when the test stimulus is shown with perceptually similar distracting stimuli. (C) Maze Learning: Average proportions of perseverations and overall errors per ‘tile’ over 22 mazes reduce when the monkey repeats the same maze. Mazes pathlength varied from 5 to 12 tiles. The result indexes spatial learning and successful error monitoring. (D) Flexible-Learning: Average learning curves (left) over seven sessions indicate slower learning (more trials to criterion performance) for extra-than intra-dimensional block switches of target features (right). (E) Effort Control: Example results from a block of 80 trials of the Effort Control task (left panel) shows the monkey chose generally more likely the option with higher reward outcome (y-axis values are above 0.5) and that his reward sensitivity increases when the absolute difference of effort between options is larger, indexing he resolves a conflict between reward and effort. Overall, the monkey preferred the lower-effort option (right panel). Sigmoidal fits provide intersection values for y=0.5 and x=0.5 to estimate reward-and effort-sensitivity. (F) Object-Sequence Learning. Learning the temporal order of objects is reflected in reduced errors, indexing successful error monitoring (left panel). The monkey choses incorrectly the order-irrelevant distractor at the temporal position at which it is similar to the correct target object for that slot, which was slot two for the initial sequence (blue line) and slot four 4 at a later encounter (red line). This indexes successful distractor interference control and serial item order learning.

### M-USE extendibility to experiments with gaze control and precise timing requirements

Multi-task performance using M-USE can incorporate tracking gaze and can be extended to set-ups that require high temporal precision and accuracy. M-USE has integrated the use of Tobii eye trackers running at 120 Hz, 300 Hz, or 600 Hz sampling frequencies. Calibration is achieved with a multi-point calibration task (1, 3, 5 or 9 symmetrically-distributed points) which is called by default at the beginning of a session when M-USE detects a connected eye tracker. The calibration task can be activated during inter-trial periods while subjects perform tasks or during the Session level’s SelectTask state, to allow re-calibration to correct for drift, etc,. Tasks can incorporate active gaze components, e.g. by requiring subjects to maintain fixation on an object for a minimum duration, or can use passive tracking to measure the distribution of fixations or information sampling behavior through saccadic eye movements and fixations (**Figure 7**).

**Figure 7.**
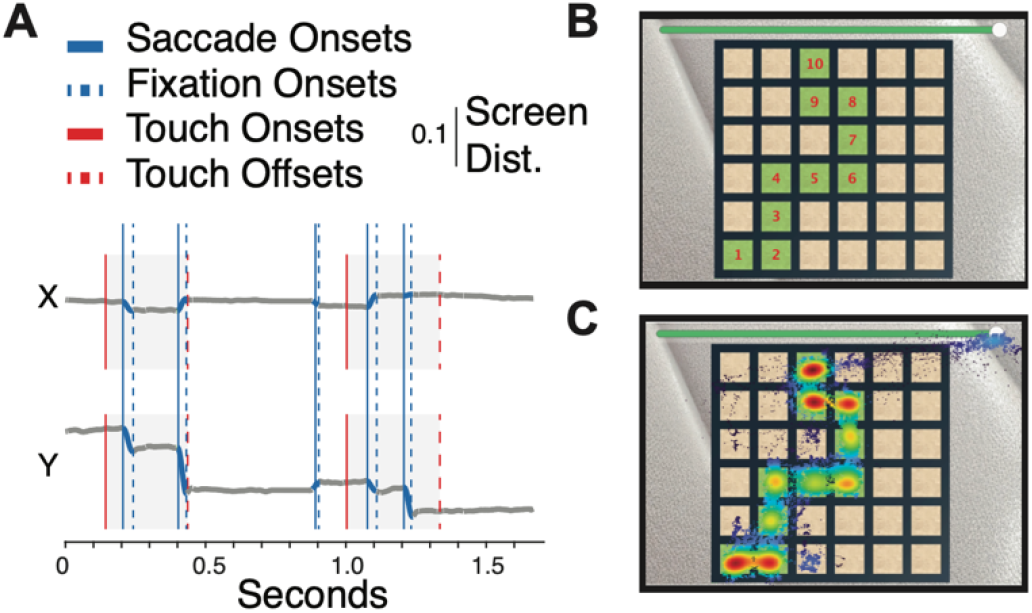
Monitoring gaze and touch during task performance. (A) Example horizontal and vertical traces of gaze (blue) and touch (red) behavior during task performance. Saccade and fixation onsets (solid and dashed vertical bars) are classified with a robust median thresholding algorithm (Voloh et al., 2019). (B) An example path of the Maze Learning task. (C) Example of a density heat map of gaze coordinates measured while recalling the trajectory shown in (B). The high gaze precision is enabled by calibrating gaze with an 9-point calibration routine built into-M-USE.

For experiments that require millisecond precise timing of events, M-USE has a configuration that flashes white and black alternating square in two corners of the subject’s monitor. The timing of the bright flashes allows reconstructing the physical temporal onset of frames and the identification of delayed or stuck frames. We have shown that frame onsets can be precisely reconstructed (**Figure 3**) with an analysis script that accompanies M-USE’s Github repository (*see* Appendix).

## 4. Discussion

We have presented M-USE, a Unity-based software platform for developing and controlling multiple cognitive tasks, and surveyed results from NHPs performing up to five tasks in daily behavioral sessions. These results demonstrate (*i*) classical pop-out and set-size effects during visual search, (*ii*) delay-dependent accuracy decline in delayed match-to-sample, (*iii*) reduced working memory performance with perceptually-similar probe and distractor stimuli, (*iv*) learning curves and set shifting switch costs, (*v*) gradual reductions of perseverative errors when learning spatial paths (Maze Learning task), and object sequences (Object Sequence learning task), and (*vi*) a motivational index derived from a sigmoidal relationship of costs (effort) and benefits (token gains) in the Effort Control task (**Figure 6**). These results illustrate that M-USE’s pre-configured tasks enable the efficient assessment of multiple cognitive and motivational performance metrics, metrics that map onto clinically meaningful cognitive constructs that encompass core constructs of the Research Domain Criteria (*RDoC*) the National Institutes of Mental Health (NIMH) considers particularly relevant for identifying the neurobiological basis of mental disorders (**Figures 4 and 5**).

### M-USE’s versatility

M-Use is a versatile software platform for the development and control of complex cognitive neuroscience experiments. Some of the key features underlying this versatility include:

- A modular structure with common elements of experiment design isolated from task development concerns in the Modules, enabling rapid prototyping and development of new tasks (**Figure 1**).
- Support for multiple tasks in single sessions, enabling efficient within-subject comparisons across multiple measures.
- Support for builds on multiple platforms, enabling use with standard computers, and with the support for web-based tasks, the ability to leverage large-scale data collection.
- Support for communication with external devices including eyetrackers, I/O devices (such as our custom Arduino-based ‘SyncBox’, *see* Appendix), neural data acquisition systems, reward delivery pumps, light-sensing diodes (for tracking frame onsets), touchscreens, or joysticks, allowing for the control of a wide range of experimental setups (**Figure 2**).
- The ability to develop and control traditional behavioral neuroscience tasks with *static* screens, but also to leverage the rich complexity of stimulus and input options required for immersive, *active* tasks (Watson et al., 2019b), while still maintaining rigorous experimental control.
- A library of pre-existing tasks that target well-known neuropsychological constructs, enabling multi-dimensional comparisons of performance on any task to existing benchmarks (**Figures 4, 5**).
- A common multidimensional stimulus space, using Quaddle objects whose 2D or 3D generation is automatized using Blender (**Figure 4**).

### The M-USE platform enables tasks with varying degrees of immersion

As a powerful video game engine, Unity can enable rich, immersive experiences. For the M-USE version released with this article, we focused on developing a Task Library that ranged across cognitive domains with relevance to neuropsychiatric disorders. Various forms of animated visual feedback provide game-like qualities of the platform, including halos around selected objects, flashing tiles in the spatial Maze Learning task, animated balloon inflation in the Effort Control task, the updating of progress bar sliders in the Maze Learning task and Object Sequence learning task, the animated token update in the other tasks. More immersive and active tasks can be implemented involving, for example, spatially navigating 3D scenes and selecting objects via joysticks (*see* Watson et al., in preparation; Watson et al., 2019b). Preliminary evidence from these immersive tasks suggests that learning complex context-dependent rules while navigating through rich environments with a joystick is governed by similar principles as learning logically-equivalent rules in more static tasks like Flex Learning (Watson et al., in preparation; see also Barrett et al., 2022). This is consistent with an understanding of the prefrontal cortex as having the ability to isolate and abstract important elements of tasks in a flexible, goal-directed manner. Having the ability to implement immersive studies promises increased ecological validity of cognitive evaluations (Kingstone et al., 2008; McColeman et al., 2020; Risko et al., 2012). The M-USE platform facilitates adding ecologically relevant features to tasks by allowing running the same state flow logic and task structure in traditional 2D static contexts, at different levels of 3D immersion, and with differing active navigational requirements. Especially when combined with M-USE’s flexible I/O connectivity, these features sets the platform apart from other experimental suites that leverage various 3D and gaming capabilities in experimental design (e.g. Bebko & Troje, 2020; Bovo et al., 2022; Brookes et al., 2018; Doucet et al., 2016; Jangraw et al., 2014; Razavi et al., 2022; Schuetz et al., 2023; Watson et al., 2019b; Cutone & Wilcox, 2021).

Other video game engines with a similar level of complexity and power as Unity include Unreal, CRYENGINE, and the open-source Godot. Unity has a large pre-existing userbase, and is taught in many university programs, making it an attractive option for students to engage with. Unity may also be simpler to use than Unreal or CRYENGINE, and more feature-rich and fully-documented than Godot, making it a logical choice to house a platform like M-USE.

### M-USE’s Task Library enables clinically meaningful cross-species cognitive profiling

M-USE was designed to facilitate cross-species evaluation of cognitive profiles and to support preclinical, translational research for improving diagnostics and treatment development for neuropsychiatric disorders (Barnett et al., 2016). The pre-configured task paradigms in M-USE serve this purpose by measuring cognitive and motivational domains that encompass the core RDoC constructs that were each previously validated as being translationally meaningful when assessed in humans and in NHPs (Friedman & Robbins, 2022; Oikonomidis et al., 2017; Palmer et al., 2021). The majority of this prior validation has been achieved using the Cambridge Automated Neuropsychological Test Associated Battery (CANTAB), which allows multi-task assessments of cognitive control, working memory, attention, relational memory and motivation. CANTAB is commercially available as a NHP version (Lafayette Instrument Company, Lafayette, IN) with tasks controlled by a client-server system (Cardinal & Aitken, 2010), and with reference performance levels available for rhesus monkeys (Weed et al., 1999). M-USE’s pre-configured tasks are variations of tasks that are realized in CANTAB or other test batteries because of their translational, clinical relevance. We illustrate this for a subset of the tasks:

- The Flex-Learning task measures set shifting abilities similar to CANTAB (Weed et al., 1999) and to the Cognitive Neuroscience Treatment Research to Improve Cognition in Schizophrenia (CNTRICS) test battery (Barch et al., 2009; Gilmour et al., 2013). It assesses both set shifting performance and reward and loss sensitivity (via variable gains and loss of tokens) in order to separate processes of rule-guided cognitive control, supported by the lateral prefrontal cortex, and valuation-based outcome recognition processes needed to identify rule shifts and reversals, supported by ventral prefrontal and orbitofrontal cortices (Dias et al., 1996; Murray & Rudebeck, 2018).
- The spatial Maze Learning task is an adoption of the Groton Maze Learning Task (GMLT), which is part of the CogState (CGS) neuropsychological test battery designed to assess rule-guided learning and error monitoring domains of the MATRICS (Measurement and Treatment Research to Improve Cognition in Schizophrenia) (Buchanan et al., 2011; Pietrzak et al., 2008).
- The Effort Control task is an extension of progressive ratio (PR) paradigms used to assess motivational processes in CANTAB (Weed et al., 1999), and of approach/avoidance conflict tasks used in NHPs (Amemori et al., 2021) and is related to tasks of the touchscreen test battery EMOTICOM (Bland et al., 2016) and the ‘Effort-Expenditure for Rewards Task’ (EefRT) (Treadway et al., 2009). The motivational constructs it measures overlap with the constructs ‘reward responsiveness’ and ‘drive’ as assessed, e.g. with the Behavioral Activation Scale (BAS) in humans, both of which are lower in humans diagnosed with major depression or with a risk to develop depression (Kasch et al., 2002; McFarland et al., 2006; Meyer et al., 1999).
- The Object sequence learning task is an extension of the Paired Associates Learning (PAL) task paradigm that evaluates relational memory in the CANTAB test battery (Barnett et al., 2016). PAL assesses relational memory in NHP’s (Taffe et al., 2002; Taffe et al., 2004), and is known to be sensitive to NMDA, GABA, ACh, and 5HT-6 antagonists, as well as to alcohol and cannabis use (Barnett et al., 2016).
- The Continuous Recognition task resembles visual working memory span and self-ordered working memory tasks developed for the CANTAB to assess the updating of working memory as well as inhibitory control in humans and NHPs (Collins et al., 1998; Walker et al., 2009). The CR task requires working memory maintenance of multiple objects, but instead of a match-to-sample decision as in the delayed match-to-sample task, the CR task requires a non-match-to sample choice, i.e. a choice of a novel object that was not seen in previous trials. The CR task resembles self-ordered WM task, because subjects self-order the objects they chose, and shares similarity to n-back tasks which require a continuous monitoring of working memory content (Petrides, 2005; Wager & Smith, 2003).

The surveyed examples illustrate some overall consensus about the cognitive processes that clinically relevant tasks should assess. However, despite this consensus, there have been practical limitations testing more than a few task paradigms in NHP. Only a small subset of NHP studies report behavioral measures from more than two tasks. For example, when assessing drugs with pro-cognitive effects (Hassani et al., 2023) or drugs relevant for tracking or treating the degree of substance use disorder, single NHP studies have combined three or four tasks in a single study including visual discrimination, reversal learning, set shifting and working memory tasks (Gould et al., 2012; Porter et al., 2011; Kangas et al., 2016). These multi-task studies contribute impressive insights into the specificity of cognitive deficits that accompany acute and chronic drug abuse (Galbo-Thomma & Czoty, 2023), but even they did not use all 3 or 4 tasks in a single behavioral assessment session, but rather split daily sessions into those assessing, e.g. working memory, and other sessions assessing reward learning and set shifting (John et al., 2018). Assessing performance of all relevant tasks in single sessions would shorten the time of data collection and reduce the variance of the behavioral results that stems from collecting data on different days. M-USE addresses these limitations by making multi-task assessments in NHP more efficient and enabling easy within-session measurement of more than a few tasks.

The breadth of cognitive domains targeted by the Task Library gives M-USE additional benefits over existing testing regimes. We routinely measure up to five tasks per session in NHP and extract per session not only behavioral metrics for executive functions like set shifting, error monitoring and working memory, but also for long-term relational memory of spatial and object associations (Maze Learning and Object Sequence learning), and motivational effort control. This illustrates that a comprehensive measurement of core cognitive and motivational domains can be captured in single behavioral sessions in NHPs, a particular worthy achievement. Such multi-dimensional cognitive/motivational profiles carry high clinical potential, as the differential diagnosis of disorders relies on the dissociation of cognitive abilities in order to infer the clinical role of the brain systems that underlie these abilities. Even common disorders like Mild Cognitive Impairment (MCI) entail varying combination of deficits: memory decline is a core deficit in MCI, but sub-types exist with additional or primary executive function deficits (Belleville et al., 2017; Prado et al., 2019). Separating these subtypes requires testing multiple cognitive functions, which serves to improve diagnostics, treatment selection and prognosis as MCI is a precursor of Alzheimer’s disease (McKhann et al., 2011). In summary, even localized and targeted pharmacological, genetic (Romberg et al., 2013), or experimental manipulations can lead to widespread changes of attention, memory and learning functions, which necessitate multi-task assessment of cognitive profiles.

#### Limitations and Future Extensions of M-USE

We introduced M-USE as a novel Unity-based platform that controls seven pre-configured tasks and that can generate standalone computer or web-based applications using these or novel tasks. Beyond the current capabilities of M-USE can be extended for future use-cases in multiple directions, several of which are actively under development or planned future extensions.

- Adaptive staircase selection of trial and block parameters, enabling an efficient determination of psychometric thresholds for subjects with varying abilities.
- Incorporation of standard psychophysical tools like fully customizable Gabor patch generation, which can solidify M-USE’s use as a psychophysical testing platform.
- Verified Linux, Mac, iOS and Android builds, further extending the breadth of M-USE.
- Verified performance in stereoscopically-rendered virtual 3D environments (e.g. using goggles)
- Incorporation of efficient video-based pose-tracking, enabling passive capture of this information for later analysis, as well as pose-or position-dependent tasks.
- The development of a standalone experimenter display app allowing a single experimenter to control multiple instances of M-USE running on different machines. Such a multi-site control will enhance the efficiency of data collection, similar to what is achieved with behavioral assessment devices used in rodents.

In summary, M-USE is a novel platform for an increasingly comprehensive and efficient assessment of cognitive and motivational skills in humans and nonhuman primates. Data collected with M-USE begin to show the potential of measuring wide-ranging cognitive profiles of individual subjects with advanced tasks and promises to support critical preclinical research needed to enhance diagnostics and treatment development for neuropsychiatric disorders.

## Appendix

M-USE is surveyed on the website http://m-use.psy.vanderbilt.edu. Files and resources are available on Github repositories:

- M-USE executables and Unity source code, as well as default resource and configuration files, are available on the main M-USE repository (https://github.com/Multitask-Unified-Suite-for-Expts).
- An *Experimenter Manual* describing how to install, run, and control M-USE experiments, a *Developer Manual* describing how to create new Tasks, and containing details of the software architecture, and providing a complete Configuration file reference are available as part of the M-USE Documentation repository (https://github.com/Multitask-Unified-Suite-for-Expts/M-USE_Documentation).
- The scripts used to generate the various files needed by M-USE, including configuration files, the multidimensional Quaddle stimuli, additional files defining the paths used by the Maze Learning task, and the context backgrounds used by most tasks are found in the M-USE Support File Generation repository (https://github.com/Multitask-Unified-Suite-for-Expts/M-USE_SupportFileGeneration).
- Matlab (The Mathworks) analysis scripts that parse M-USE session data folders, unite each data type (Frame, Trial, etc) into a corresponding.mat object, and perform various preliminary analyses (including timing analyses) are found in the M-USE Analysis repository (https://github.com/Multitask-Unified-Suite-for-Expts/M-USE_Analysis).
- Documentation of the Synchbox and hardware (LED light sensor mounts) that we used for time reconstruction is online available at https://github.com/att-circ-contrl/SynchBox.

## Financial Disclosures

The authors declare no competing financial interests.

## Acknowledgements

This work was supported by the National Institute of Mental Health of the National Institutes of Health under Award Number R01MH129641 (TW). The content is solely the responsibility of the authors and does not necessarily represent the official views of the National Institutes of Health.

## Notes

http://m-use.psy.vanderbilt.edu

### Competing Interest Statement

The authors have declared no competing interest.

## References

1. Altschul, D., Jensen, G., & Terrace, H. (2017). Perceptual category learning of photographic and painterly stimuli in rhesus macaques (Macaca mulatta) and humans. PLOS ONE, 12(9). 10.1371/journal.pone.0185576

2. Amemori, K.-I., & Graybiel, A. M. (2012). Localized microstimulation of primate pregenual cingulate cortex induces negative decision-making. Nature Neuroscience, 15(5), 776–785. 10.1038/nn.3088

3. Amemori, S., Graybiel, A. M., & Amemori, K.-I. (2021). Causal Evidence for Induction of Pessimistic Decision-Making in Primates by the Network of Frontal Cortex and Striosomes. Frontiers in Neuroscience, 15(107). 10.3389/fnins.2021.649167

4. Aragona, M. (2014). Epistemological reflections about the crisis of the DSM-5 and the revolutionary potential of the RDoC project. *Dialogues in Philosophy*, Mental and Neuro Sciences, 7(1), 11–20.

5. Axelsson, S. F. A., Horst, N. K., Horiguchi, N., Roberts, C.A., & Robbins, T.W. (2021). Flexible versus Fixed Spatial Self-Ordered Response Sequencing: Effects of Inactivation and Neurochemical Modulation of Ventrolateral Prefrontal Cortex. Journal of Neuroscience, 41(34), 7246–7258. 10.1523/JNEUROSCI.0227-21.2021

6. Badcock, J. C., Michie, P. T., & Rock, D. (2005). Spatial working memory and planning ability: Contrasts between schizophrenia and bipolar I disorder. Cortex, 41(6), 753–763. 10.1016/S0010-9452(08)70294-6

7. Bakker, G., Vingerhoets, C., Bloemen, O. J. N. A., Sahakian, B. J., Booij, J., Caan, M. W. A., van Amelsvoort, T. A. M. J., &. (2020). The muscarinic M-1 receptor modulates associative learning and memory in psychotic disorders. NEUROIMAGE-CLINICAL NeuroImage-Clin., 27(45). 10.1016/j.nicl.2020.102278

8. Ball, K., Edwards, J. D., & Ross, L. A. (2007). The impact of speed of processing training on cognitive and everyday functions. Journals of Gerontology Series B-Psychological Sciences and Social Sciences, 62(1), 19–31. 10.1093/geronb/62.special\_issue\_1.19

9. Barch, D. M., Braver, T. S., Carter, C. S. A., Poldrack, R. A., & Robbins, T. W. (2009). CNTRICS Final Task Selection: Executive Control. Schizophrenia Bulletin, 35(1), 115–135. 10.1093/schbul/sbn154

10. Barnett, J. H., Blackwell, A. D., Sahakian, B. J., & Robbins, T. W. (2016). The Paired Associates Learning (PAL) Test: 30 Years of CANTAB Translational Neuroscience from Laboratory to Bedside in Dementia Research. Current Topics in Behavioral Neurosciences, 28(93), 449–474. 10.1007/7854\_2015\_5001

11. Barrett, R. C. A., Poe, R., O’Camb, J. W., Woodruff, C., Harrison, S. M., Dolguikh, K., Chuong, C., Klassen, A. D., Zhang, R., Joseph, R. B., & Blair, M. R. (2022). Comparing virtual reality, desktop-based 3D, and 2D versions of a category learning experiment. PLoS One, 17(10), e0275119. 10.1371/journal.pone.0275119

12. Bebko, A. O., & Troje, N. F. (2020). bmlTUX: Design and Control of Experiments in Virtual Reality and Beyond. iPerception, 11(4), 2041669520938400. 10.1177/2041669520938400

13. Belleville, S., Fouquet, C., Hudon, C., Zomahoun, Vignon, H. T., Croteau, J., & Quebec, C. F. T. E. I. O. A.D. (2017). Neuropsychological Measures that Predict Progression from Mild Cognitive Impairment to Alzheimer’s type dementia in Older Adults: a Systematic Review and Meta-Analysis. Neuropsychology Review, *27*(4, SI), 328-353. 10.1007/s11065-017-9361-5

14. Berger, M., Calapai, A., Stephan, V., Niessing, M. A., Burchardt, L., Gail, A., & Treue, S. (2018). Standardized automated training of rhesus monkeys for neuroscience research in their housing environment. Journal of Neurophysiology, 119(3), 796–807. 10.1152/jn.00614.2017

15. Berwian, I. M., Wenzel, J. G., Collins, A. G. E., Seifritz, E., Stephan, K. E., Walter, H., & Huys, Q. J. M. (2020). Computational Mechanisms of Effort and Reward Decisions in Patients With Depression and Their Association With Relapse After Antidepressant Discontinuation. JAMA Psychiatry, 77(5), 513–522. 10.1001/jamapsychiatry.2019.4971

16. Bornert, P., & Bouret, S. (2021). Locus coeruleus neurons encode the subjective difficulty of triggering and executing actions. PLOS Biology, 19(12). 10.1371/journal.pbio.3001487

17. Boroujeni, K. B., Watson, M., & Womelsdorf, T. (2022). Gains and Losses Affect Learning Differentially at Low and High Attentional Load. J Cogn Neurosci, 34(10), 1952–1971. 10.1162/jocn_a_01885

18. Bovo, R., Giunchi, D., Steed, A., & Heinis, T. (2022). MR-RIEW: An MR Toolkit for Designing Remote Immersive Experiment Workflows. In. IEEE. 10.1109/vrw55335.2022.00234

19. Brookes, J., Warburton, M., Alghadier, M., Mon-Williams, M. A., & Mushtaq, F. (2018). Studying human behaviour with virtual reality: The Unity Experiment Framework. biorxiv. 10.1101/459339

20. Brown, V. J., & Tait, D. S. (2016). Attentional set-shifting across species. In T. W. Robbins & B. J. Sahakian (Eds.), Translational Neuropsychopharmacology (Vol. 28, pp. 363–395). Springer International Publishing.

21. Buchanan, R. W., Keefe, R. S. E., Umbricht, D. A., Green, M. F., Laughren, T., & Marder, S. R. (2011). The FDA-NIMH-MATRICS Guidelines for Clinical Trial Design of Cognitive-Enhancing Drugs: What Do We Know 5 Years Later? Schizophrenia Bulletin, 37(6), 1209–1217. 10.1093/schbul/sbq038

22. Buckley, M. J., Mansouri, F. A., Hoda, H., Mahboubi, Majid, Browning, P. G. F., Kwok, S. C., Phillips, A. A., & Tanaka, K. (2009). Dissociable Components of Rule-Guided Behavior Depend on Distinct Medial and Prefrontal Regions. Science, 325(5936), 52–58. 10.1126/science.1172377

23. Buzsaki, G., & Tingley, D. (2018). Space and Time: The Hippocampus as a Sequence Generator. Trends in Cognitive Sciences, *22*(10, SI), 853-869. 10.1016/j.tics.2018.07.006

24. Calapai, A., Berger, M., Niessing, M., Heisig, K. A., Brockhausen, R., Treue, S., & Gail, A. (2017). A cage-based training, cognitive testing and enrichment system optimized for rhesus macaques in neuroscience research. Behavior Research Methods, 49(1), 35–45. 10.3758/s13428-016-0707-3

25. Callahan, M. J. (1999). Combining tacrine with milameline reverses a scopolamine-induced impairment of continuous performance in rhesus monkeys. Psychopharmacology, 144(3), 234–238. 10.1007/s002130050998

26. Cardinal, R. N., & Aitken, M. R. F. (2010). Whisker: A client-server high-performance multimedia research control system. Behavior Research Methods, 42(4), 1059–1071. 10.3758/BRM.42.4.1059

27. Champod, A. S., & Petrides, M. (2007). Dissociable roles of the posterior parietal and the prefrontal cortex in manipulation and monitoring processes. Proceedings of the National Academy of Sciences of the United States of America, 104(37), 14837–14842. 10.1073/pnas.0607101104

28. Chase, H. W., Clark, L., Sahakian, B. J., Bullmore, T., E., & Robbins, T.W. (2008). Dissociable roles of prefrontal subregions in self-ordered working memory performance. Neuropsychologia, 46(11), 2650–2661. 10.1016/j.neuropsychologia.2008.04.021

29. Cocker, P. J., Hosking, J. G., Benoit, J., Winstanley, & A., C. (2012). Sensitivity to Cognitive Effort Mediates Psychostimulant Effects on a Novel Rodent Cost/Benefit Decision-Making Task. Neuropsychopharmacology, 37(8), 1825–1837. 10.1038/npp.2012.30

30. Collins, A. G., Cavanagh, J. F., & Frank, M. J. (2014). Human EEG uncovers latent generalizable rule structure during learning. J Neurosci, 34(13), 4677–4685. 10.1523/JNEUROSCI.3900-13.2014

31. Collins, A. G. E., & Frank, M. J. (2013). Cognitive control over learning: Creating, clustering, and generalizing task-set structure. Psychological Review, 120(1), 190–229.

32. Collins, P., Roberts, A. C., Dias, R., Everitt, B. J., & Robbins, T. W. (1998). Perseveration and strategy in a novel spatial self-ordered sequencing task for nonhuman primates: Effects of excitotoxic lesions and dopamine depletions of the prefrontal cortex. Journal of Cognitive Neuroscience, 10(3), 332–354. 10.1162/089892998562771

33. Cuthbert, B. N., & Insel, T. R. (2013). Toward the future of psychiatric diagnosis: the seven pillars of RDoC. BMC Med, 11, 126. 10.1186/1741-7015-11-126

34. Cutone, M. D., & Wilcox, L. M. (2021). PsychXR. Available from https://github.com/mdcutone/psychxr.

35. Der-Avakian, A., Barnes, S. A., Markou, A. A., & Pizzagalli, D. A. (2016). Translational Assessment of Reward and Motivational Deficits in Psychiatric Disorders. Current Topics in Behavioral Neurosciences, 28(195), 231–262. 10.1007/7854\_2015\_5004

36. Dias, R., Robbins, T. W., & Roberts, A. C. (1996). Dissociation in prefrontal cortex of affective and attentional shifts. Nature, 380(6569), 69–72. 10.1038/380069a0

37. Doucet, G., Gulli, R. A., & Martinez-Trujillo, J. C. (2016). Cross-species 3D virtual reality toolbox for visual and cognitive experiments. J Neurosci Methods, 266, 84–93. 10.1016/j.jneumeth.2016.03.009

38. Dowson, J. H., McLean, A., Bazanis, E., Toone, B., Young, S. A., Robbins, T. W., & Sahakian, B. J. (2004). Impaired spatial working memory in adults with attention-deficit/hyperactivity disorder: comparisons with performance in adults with borderline personality disorder and in control subjects. Acta Psychiatrica Scandinavica, 110(1), 45–54. 10.1111/j.1600-0447.2004.00292.x

39. Edwards, J. D., Wadley, V. G., Myers, R. S., Roenker, D. L., Cissell, G. M., & Ball, K. K. (2002). Transfer of a speed of processing intervention to near and far cognitive functions. GERONTOLOGY, 48(5), 329–340. 10.1159/000065259

40. Fray, P. J., Robbins, T. W., & Sahakian, B. J. (1996). Neuropsychiatric applications of CANTAB. International Journal of Geriatric Psychiatry, 11(4), 329–336.

41. Fried, R., Hirshfeld-Becker, D., Petty, C. A., Batchelder, H., & Biederman, J. (2015). How Informative Is the CANTAB to Assess Executive Functioning in Children With ADHD? A Controlled Study. Journal of Attention Disorders, 19(6, SI), 468–475. 10.1177/1087054712457038

42. Friedman, N. P., & Robbins, T. W. (2022). The role of prefrontal cortex in cognitive control and executive function. Neuropsychopharmacology, 47(1), 72–89. 10.1038/s41386-021-01132-0

43. Friedman, N. P., Miyake, A., Corley, R. P., Young, S. E., DeFries, J. C., & and Hewitt, J. K. (2006). Not all executive functions are related to intelligence. Psychological Science, 17(2), 172–179. 10.1111/j.1467-9280.2006.01681.x

44. Galbo-Thomma, L. K., & Czoty, P. W. (2023). The Use of Touchscreen-Based Methods to Characterize Effects of Psychoactive Drugs on Executive Function in Nonhuman Primates. Current Pharmacology Reports. 10.1007/s40495-023-00337-9

45. Gilmour, G., Arguello, A., Bari, A., Brown, V. J., Carter, C., Floresco, S. B., Jentsch, D. J., Tait, D. S., Young, J. W., & Robbins, T. W. (2013). Measuring the construct of executive control in schizophrenia: Defining and validating translational animal paradigms for discovery research. Neuroscience and Biobehavioral Reviews, *37*(9, B), 2125-2140. 10.1016/j.neubiorev.2012.04.006

46. Gold, J. M., Strauss, G. P., Waltz, J. A., Robinson, B. M., Brown, J. K., & Frank, M. J. (2013). Negative symptoms of schizophrenia are associated with abnormal effort-cost computations. Biol Psychiatry, 74(2), 130–136. 10.1016/j.biopsych.2012.12.022

47. Gould, R. W., Gage, H. D., & Nader, M. A. (2012). Effects of chronic cocaine self-administration on cognition and cerebral glucose utilization in Rhesus monkeys. Biol Psychiatry, 72(10), 856–863. 10.1016/j.biopsych.2012.05.001

48. Harlow, H. F. (1949). The formation of learning sets. Psychological Review, 56(1), 51–65. 10.1037/h0062474

49. Harvey, P. D. (2023). Cognition in Schizophrenia: MATRICS Consensus Cognitive Battery (MCCB): Development, Characteristics, and Usage. In G. J. Boyle, Y. Stern, D. J. Stein, B. J. Sahakian, C. J. Golden, T. M.-C. Lee, & S.-H. A. Chen (Eds.), The SAGE Handbook of Clinical Neuropsychology: Clinical Neuropsychological Assessment and Diagnosis (p. 617). SAGE Publications, Ltd.

50. Hassani, S. A., Lendor, S., Neumann, A., Sinha Roy, K., Banaie Boroujeni, K., Hoffman, K. L., Pawliszyn, J., & Womelsdorf, T. (2021). Dose-Dependent Dissociation of Pro-cognitive Effects of Donepezil on Attention and Cognitive Flexibility in Rhesus Monkeys. Biol Psychiatry Glob Open Sci, 3(1), 68–77. 10.1016/j.bpsgos.2021.11.012

51. Hassani, S. A., Neumann, A., Russell, J., Jones, C. K., & Womelsdorf, T. (2023). M1-selective muscarinic allosteric modulation enhances cognitive flexibility and effective salience in nonhuman primates. Proc Natl Acad Sci U S A, 120(18), e2216792120. 10.1073/pnas.2216792120

52. Heuer, E., & Bachevalier, J. (2013). Working memory for temporal order is impaired after selective neonatal hippocampal lesions in adult rhesus macaques. Behav. Brain Res., 239(49), 55–62. 10.1016/j.bbr.2012.10.043

53. Jang, A. I., Costa, V. D., Rudebeck, P. H. A., Chudasama, Y., Murray, E. A., & Averbeck, B. B. (2015). The Role of Frontal Cortical and Medial-Temporal Lobe Brain Areas in Learning a Bayesian Prior Belief on Reversals. Journal of Neuroscience, 35(33), 11751–11760. 10.1523/JNEUROSCI.1594-15.2015

54. Jangraw, D. C., Johri, A., Gribetz, M., & Sajda, P. (2014). NEDE: an open-source scripting suite for developing experiments in 3D virtual environments. J Neurosci Methods, 235, 245–251. 10.1016/j.jneumeth.2014.06.033

55. Joyce, E. M., Hutton, S. B., Mutsatsa, S. H., & Barnes, T. R. E. (2005). Cognitive heterogeneity in first-episode schizophrenia. British Journal of Psychiatry, 187(55), 516–522. 10.1192/bjp.187.6.516

56. Kangas, B. D., Leonard, M. Z., Shukla, V. G., Alapafuja, S. O., Nikas, S. P., Makriyannis, A., & Bergman, J. (2016). Comparisons of Δ9-Tetrahydrocannabinol and Anandamide on a Battery of Cognition-Related Behavior in Nonhuman Primates. J Pharmacol Exp Ther, 357(1), 125–133. 10.1124/jpet.115.228189

57. Kangas, B. D., & Bergman, J. (2017). Touchscreen technology in the study of cognition-related behavior. Behavioural Pharmacology, 28(8), 623–629. 10.1097/FBP.0000000000000356

58. Kasch, K. L., Rottenberg, J., Arnow, B. A., & Gotlib, I. H. (2002). Behavioral activation and inhibition systems and the severity and course of depression. Journal of Abnormal Psychology, 111(4), 589–597. 10.1037//0021-843X.111.4.589

59. Keller, E. L., Lee, K.-M., Park, S.-W., & Hill, J. A. (2008). Effect of Inactivation of the Cortical Frontal Eye Field on Saccades Generated in a Choice Response Paradigm. Journal of Neurophysiology, 100(5), 2726–2737. 10.1152/jn.90673.2008

60. Kennedy, B. L., & Mather, M. (2019). Neural Mechanisms Underlying Age-Related Changes in Attentional Selectivity. In G. R. Samanez-Larkin (Ed.), The aging brain:functional adaptation across adulthood The aging brain:functional adaptation across adulthood (pp. 45–72). American Psychological Association.

61. Kingstone, A., Smilek, D., & Eastwood, J. D. (2008). Cognitive Ethology: A new approach for studying human cognition. British Journal of Psychology, 99(3), 317–340. 10.1348/000712607X251243

62. Lange, H. S., Cannon, C. E., Drott, J. T. A., Kuduk, S. D., & Uslaner, J. M. (2015). The M1 Muscarinic Positive Allosteric Modulator PQCA Improves Performance on Translatable Tests of Memory and Attention in Rhesus Monkeys. Journal of Pharmacology and Experimental Therapeutics, 355(3), 442–450. 10.1124/jpet.115.226712

63. Langley, C., Sahakian, B. J., & Robbins, T. W. (2023). Cambridge Neuropsychological Test Automated Battery (CANTAB). In G. J. Boyle, Y. Stern, D. J. Stein, B. J. Sahakian, C. J. Golden, T. M.-C. Lee, & S.-H. A. Chen (Eds.), The SAGE Handbook of Clinical Neuropsychology: Clinical Neuropsychological Assessment and Diagnosis (p. 435). SAGE Publications, Ltd.

64. Levy, I., Snell, J., Nelson, A. J., Rustichini, A. A., & Glimcher, P. W. (2010). Neural Representation of Subjective Value Under Risk and Ambiguity. Journal of Neurophysiology, 103(2), 1036–1047. 10.1152/jn.00853.2009

65. Lieberman, J. A., Dunbar, G., Segreti, A. C. A., Girgis, R. R., Seoane, F., Beaver, J. S., Duan, Naihua, & Hosford, D. A. (2013). A Randomized Exploratory Trial of an Alpha-7 Nicotinic Receptor Agonist (TC-5619) for Cognitive Enhancement in Schizophrenia. Neuropsychopharmacology, 38(6), 968–975. 10.1038/npp.2012.259

66. McColeman, C., Thompson, J., Anvari, N., Azmand, S. J., Barnes, J., Barrett, R. C. A., Byliris, R., Chen, Y., Dolguikh, K., Fischler, K., Harrison, S., Hayre, R. S., Poe, R., Swanson, L., Tracey, T., Volkanov, A., Woodruff, C., Zhang, R., & Blair, M. (2020). Digit eyes: Learning-related changes in information access in a computer game parallel those of oculomotor attention in laboratory studies. Atten Percept Psychophys, 82(5), 2434–2447. 10.3758/s13414-020-02019-w

67. McFarland, B. R., Shankman, S. A., Tenke, C. E., Bruder, G. E., & Klein, D. N. (2006). Behavioral activation system deficits predict the six-month course of depression. Journal of Affective Disorders, 91(2-3), 229–234. 10.1016/j.jad.2006.01.012

68. McKhann, G. M., Knopman, D. S., Chertkow, H., Hyman, B. T., Jack, C. R., Kawas, C. H., Klunk, W. E., Koroshetz, W. J., Manly, J. J., Mayeux, R., Mohs, R. C., Morris, J. C., Rossor, M. N., Scheltens, P., Carrillo, M. C., Thies, B., Weintraub, S., & Phelps, C. H. (2011). The diagnosis of dementia due to Alzheimer’s disease: recommendations from the National Institute on Aging-Alzheimer’s Association workgroups on diagnostic guidelines for Alzheimer’s disease. Alzheimers Dement, 7(3), 263–269. 10.1016/j.jalz.2011.03.005

69. Meier, M. E., & Kane, M. J. (2013). Working Memory Capacity and Stroop Interference: Global Versus Local Indices of Executive Control. Journal of Experimental Psychology-Learning Memory and Cognition, 39(3), 748–759. 10.1037/a0029200

70. Meyer, B., Johnson, S. L., & Carver, C. S. (1999). Exploring behavioral activation and inhibition sensitivities among college students at risk for bipolar spectrum symptomatology. Journal of Psychopathology and Behavioral Assessment, 21(4), 275–292. 10.1023/A:1022119414440

71. Millan, M. J., Agid, Y., Bruene, M., Bullmore, E. T., Carter, C. S., Clayton, N. S., Connor, R., Davis, S., Deakin, B., DeRubeis, R. J., Dubois, B., Geyer, M. A., Goodwin, G. M., Gorwood, P., Jay, T. M., Joels, M., Mansuy, I. M., Meyer-Lindenberg, A., Murphy, D.,... Young, L.J. (2012). Cognitive dysfunction in psychiatric disorders: characteristics, causes and the quest for improved therapy. Nature Reviews Drug Discovery, 11(2), 141–168. 10.1038/nrd3628

72. Miyake, A., Friedman, N. P., Emerson, M. J., Witzki, A. H., Howerter, A, & Wager, T. D. (2000). The unity and diversity of executive functions and their contributions to complex”frontal lobe tasks: A latent variable analysis. Cognitive Psychology, 41(1), 49–100. 10.1006/cogp.1999.0734

73. Moore, H., Geyer, M. A., Carter, C. S., & Barch, D. M. (2013). Harnessing cognitive neuroscience to develop new treatments for improving cognition in schizophrenia: CNTRICS selected cognitive paradigms for animal models. Neuroscience and Biobehavioral Reviews, 37(9, B), 2087-2091. 10.1016/j.neubiorev.2013.09.011

74. Moore, T. L., Killiany, R. J., Herndon, J. G., Rosene, D. L., & Moss, M. B. (2003). Impairment in abstraction and set shifting in aged Rhesus monkeys. *NEUROBIOLOGY OF AGING Neurobiol*. Aging, 24(1), 125–134. 10.1016/S0197-4580(02)00054-4

75. Moore, T. L., Schettler, S. P., Killiany, R. J., Herndon, J. G., Luebke, J. I., Moss, M. B., & Rosene, D. L. (2005). Cognitive impairment in aged rhesus monkeys associated with monoamine receptors in the prefrontal cortex. Behavioural Brain Research, 160(2), 208–221. 10.1016/j.bbr.2004.12.003

76. Morris, S. E., & Cuthbert, B. N. (2012). Research Domain Criteria: cognitive systems, neural circuits, and dimensions of behavior. Dialogues Clin Neurosci, 14(1), 29–37. 10.31887/DCNS.2012.14.1/smorris

77. Murray, E. A., & Rudebeck, P. H. (2018). Specializations for reward-guided decision-making in the primate ventral prefrontal cortex. Nature Reviews Neuroscience, 19(7), 404–417. 10.1038/s41583-018-0013-4

78. Mushiake, H., Saito, N., Sakamoto, K., Sato, Y., & Tanji, J. (2001). Visually based path-planning by Japanese monkeys. Cognitive Brain Research, 11(1), 165–169. 10.1016/S0926-6410(00)00067-7

79. Mushiake, H., Saito, N., Sakamoto, K., Itoyama, Yasuto, Tanji, J., &. (2006). Activity in the lateral prefrontal cortex reflects multiple steps of future events in action plans. Neuron, 50(4), 631–641. 10.1016/j.neuron.2006.03.045

80. O’Neill, J., Siembieda, D. W., Crawford, K. C., Halgren, E., Fisher, A., & Fitten, L. J. (2003). Reduction in distractibility with AF102B and THA in the macaque. Pharmacology Biochemistry and Behavior, 76(2), 301–306. 10.1016/j.pbb.2003.08.006

81. Oikonomidis, L., Santangelo, A. M., Shiba, Y. A., Clarke, F. H., Robbins, T. W., & Roberts, A. C. (2017). A dimensional approach to modeling symptoms of neuropsychiatric disorders in the marmoset monkey. Developmental Neurobiology, 77(3), 328–353. 10.1002/dneu.22446

82. Orlov, T., Yakovlev, V., Hochstein, S., & Zohary, E. (2000). Macaque monkeys categorize images by their ordinal number. Nature, 404(6773), 77–80. 10.1038/35003571

83. Palmer, D., Dumont, J. R., Dexter, T. D., Prado, M. A. M., Finger, E., Bussey, T. J., & Saksida, L. M. (2021). Touchscreen cognitive testing: Cross-species translation and co-clinical trials in neurodegenerative and neuropsychiatric disease. Neurobiology of Learning and Memory, 182(253). 10.1016/j.nlm.2021.107443

84. Pantelis, C., Barber, F. Z., Barnes, T. R. E., Nelson, H. E., & Owen, A. M. R., TW. (1999). Comparison of set-shifting ability in patients with chronic schizophrenia and frontal lobe damage. Schizophrenia Research, 37(3), 251–270. 10.1016/S0920-9964(98)00156-X

85. Passingham, R. (2021). Understanding the prefrontal cortex: selective advantage, connectivity, and neural operations. Oxford University Press.

86. Passingham, R. E., & Wise, S. P. (2012). The neurobiology of the prefrontal cortex: anatomy, evolution, and the origin of insight. Oxford University Press.

87. Perdue, B. M., Beran, M. J., & Washburn, D. A. (2018). A computerized testing system for primates: Cognition, welfare, and the Rumbaughx. Behavioural Processes, 156(SI), 37–50. 10.1016/j.beproc.2017.12.019

88. Petrides, M. (1991). Functional specialization within the dorsolateral frontal-cortex for serial order memory. Proceedings of the Royal Society B-Biological Sciences, 246(1317), 299–306. 10.1098/rspb.1991.0158

89. Petrides, M. (2005). Lateral prefrontal cortex: architectonic and functional organization. Philosophical Transactions of the Royal Society B-Biological Sciences, 360(1456), 781–795. 10.1098/rstb.2005.1631

90. Pietrzak, R. H., Maruff, P., Mayes, L. C., Roman, A.S., Sosa, J. A., & Snyder, P.J. (2008). An examination of the construct validity and factor structure of the Groton Maze Learning Test, a new measure of spatial working memory, learning efficiency, and error monitoring. Archives of Clinical Neuropsychology, 23(4), 433–445. 10.1016/j.acn.2008.03.002

91. Pietrzak, R. H., Olver, J., Norman, T., Piskulic, Danijela, Maruff, P., & Snyder, P. J. (2009). A comparison of the CogState Schizophrenia Battery and the Measurement and Treatment Research to Improve Cognition in Schizophrenia (MATRICS) Battery in assessing cognitive impairment in chronic schizophrenia. Journal of Clinical and Experimental Neuropsychology, 31(7), 848–859. 10.1080/13803390802592458

92. Pietrzak, R. H., Snyder, P. J., & Maruff, P. (2010). Amphetamine-related improvement in executive function in patients with chronic schizophrenia is modulated by practice effects. Schizophrenia Research, 124(1-3), 176–182. 10.1016/j.schres.2010.09.012

93. Pirkola, T., Tuulio-Henriksson, A., Glahn, D., Kieseppa, T. A., Haukka, J., Kaprio, J., Lonnqvist, J., & Cannon, T. D. (2005). Spatial working memory function in twins with schizophrenia and bipolar disorder. Biological Psychiatry, 58(12), 930–936. 10.1016/j.biopsych.2005.05.041

94. Poole, B. J., & Kane, M. J. (2009). Working-memory capacity predicts the executive control of visual search among distractors: The influences of sustained and selective attention. Quarterly Journal of Experimental Psychology, 62(7), 1430–1454. 10.1080/17470210802479329

95. Popiolek, M., Mandelblat-Cerf, Y., Young, D., Garst-Orozco, J., Lotarski, S. M., Stark, E., Kramer, M., Butler, C. R., & Kozak, R. (2019). In Vivo Modulation of Hippocampal Excitability by M4 Muscarinic Acetylcholine Receptor Activator: Implications for Treatment of Alzheimer’s Disease and Schizophrenic Patients. ACS Chem Neurosci, 10(3), 1091–1098. 10.1021/acschemneuro.8b00496

96. Porter, J. N., Olsen, A. S., Gurnsey, K., Dugan, B. P., Jedema, H. P., & Bradberry, C. W. (2011). Chronic cocaine self-administration in rhesus monkeys: impact on associative learning, cognitive control, and working memory. J Neurosci, 31(13), 4926–4934. 10.1523/JNEUROSCI.5426-10.2011

97. Prado, C. E., Watt, S., Treeby, M. S., Crowe, & F., S. (2019). Performance on Neuropsychological Assessment and Progression to Dementia: A Meta-Analysis. Psychology and Aging, 34(7), 954–977. 10.1037/pag0000410

98. Razavi, M., Janfaza, V., Yamauchi, T., Leontyev, A., Longmire-Monford, S., & Orr, J. (2022). OpenSync: An open-source platform for synchronizing multiple measures in neuroscience experiments. J Neurosci Methods, 369, 109458. 10.1016/j.jneumeth.2021.109458

99. Redish, A. D., Kepecs, A., Anderson, L. M., Calvin, L.O., Grissom, N. M., Haynos, A. F., Heilbronner, R.S., Herman, A. B., Jacob, S., Ma, S. A., Vilares, I., Vinogradov, S., Walters, C. J., Widge, S.A., Zick, J. L., & Zilverstand, A. (2022). Computational validity: using computation to translate behaviours across species. Philosophical Transactions of the Royal Society B-Biological Sciences, 377(1844). 10.1098/rstb.2020.0525

100. Risko, E. F., Laidlaw, K., Freeth, M., Foulsham, T., & Kingstone, A. (2012). Social attention with real versus reel stimuli: toward an empirical approach to concerns about ecological validity. Front Hum Neurosci, 6, 143. 10.3389/fnhum.2012.00143

101. Roberts, A. C., Robbins, T. W., & Everitt, B. J. (1988). The effects of intradimensional and extradimensional shifts on visual-discrimination learning in humans and non-human primates. Quarterly Journal of Experimental Psychology Section B-Comparative and Physiological Psychology, 40(4), 321–341.

102. Roelfsema, P. R., & Treue, S. (2014). Basic Neuroscience Research with Nonhuman Primates: A Small but Indispensable Component of Biomedical Research. Neuron, 82(6), 1200–1204. 10.1016/j.neuron.2014.06.003

103. Romberg, C., Horner, A. E., Bussey, T. J., & Saksida, L. M. (2013). A touch screen-automated cognitive test battery reveals impaired attention, memory abnormalities, and increased response inhibition in the TgCRND8 mouse model of Alzheimer’s disease. Neurobiology of Aging, 34(3), 731–744. 10.1016/j.neurobiolaging.2012.08.006

104. Rossi, A. F., Bichot, N. P., Desimone, R. A., & Ungerleider, L. G. (2007). Top-down attentional deficits in macaques with lesions of lateral prefrontal cortex. Journal of Neuroscience, 27(42), 11306–11314. 10.1523/JNEUROSCI.2939-07.2007

105. Rudebeck, P. H., Saunders, R. C., Lundgren, D. A. A., & Murray, E. A. (2017). Specialized Representations of Value in the Orbital and Ventrolateral Prefrontal Cortex: Desirability versus Availability of Outcomes. Neuron, 95(5), 1208+. 10.1016/j.neuron.2017.07.042

106. Sahakian, B. J., & Owen, A. M. (1992). Computerized Assessment in Neuropsychiatry Using Cantab-Discussion Paper. Journal of the Royal Society of Medicine, 85(7), 399–402.

107. Saito, N., Mushiake, H., Sakamoto, K., Itoyama, Y., & Tanji, J. (2005). Representation of immediate and final behavioral goals in the monkey prefrontal cortex during an instructed delay period. Cerebral Cortex, 15(10), 1535–1546. 10.1093/cercor/bhi032

108. Salamone, J. D., Correa, M., Farrar, A., & Mingote, S. M. (2007). Effort-related functions of nucleus accumbens dopamine and associated forebrain circuits. Psychopharmacology, 191(3), 461–482. 10.1007/s00213-006-0668-9

109. Salamone, J. D., Cousins, M. S., & Snyder, B. J. (1997). Behavioral functions of nucleus accumbens dopamine: Empirical and conceptual problems with the anhedonia hypothesis. Neuroscience and Biobehavioral Reviews, 21(3), 341–359. 10.1016/S0149-7634(96)00017-6

110. Salamone, J. D., Correa, M., Farrar, A. M., Nunes, J.E., & Pardo, M. (2009). Dopamine, behavioral economics, and effort. Frontiers in Behavioral Neuroscience, 3(173). 10.3389/neuro.08.013.2009

111. Schuetz, I., Karimpur, H., & Fiehler, K. (2023). vexptoolbox: A software toolbox for human behavior studies using the Vizard virtual reality platform. Behav Res Methods, 55(2), 570–582. 10.3758/s13428-022-01831-6

112. Schwarz, R. D., Callahan, M. J., Coughenour, L. L., Dickerson, M. R. A., Kinsora, J. J., Lipinski, W. J., Raby, C. A., Spencer, C. J., & Tecle, H. (1999). Milameline (CI-979/RU35926): A muscarinic receptor agonist with cognition-activating properties: Biochemical and in vivo characterization. Journal of Pharmacology and Experimental Therapeutics, 291(2), 812–822.

113. Scott, J. T., & Bourne, J. A. (2022). Modelling behaviors relevant to brain disorders in the nonhuman primate: Are we there yet? Progress in Neurobiology, 208(361). 10.1016/j.pneurobio.2021.102183

114. Snyder, P. J., Jackson, C. E., Piskulic, D. A., Olver, J., Norman, T., & Maruff, P. (2008). Spatial working memory and problem solving in schizophrenia: The effect of symptom stabilization with atypical antipsychotic medication. Psychiatry Research, 160(3), 316–326. 10.1016/j.psychres.2007.07.011

115. Taffe, M. A., Weed, M. R., Gutierrez, T., Davis, S. A., & Gold, L. H. (2002). Differential muscarinic and NMDA contributions to visuo-spatial paired-associate learning in rhesus monkeys. Psychopharmacology, 160(3), 253–262. 10.1007/s00213-001-0954-5

116. Taffe, M. A., Weed, M. R., Gutierrez, T., Davis, S. A., & Gold, L. H. (2004). Modeling a task that is sensitive to dementia of the Alzheimer’s type: individual differences in acquisition of a visuo-spatial paired-associate learning task in rhesus monkeys. Behavioural Brain Research, 149(2), 123–133. 10.1016/S0166-4328(03)00214-6

117. Terrace, H. S. (2005). The simultaneous chain: a new approach to serial learning. Trends in Cognitive Science, 9(4), 202–210. 10.1016/j.tics.2005.02.003

118. Treadway, M. T., Buckholtz, J. W., Schwartzman, A. N., and Lambert, W. E., & Zald, D. H. (2009). Worth the‘EEfRT’? The Effort Expenditure for Rewards Task as an Objective Measure of Motivation and Anhedonia. PLOS ONE, 4(8). 10.1371/journal.pone.0006598

119. Unsworth, N., & Robison, M. K. (2017). The Importance of Arousal for Variation in Working Memory Capacity and Attention Control: A Latent Variable Pupillometry Study. Journal of Experimental Psychology-Learning Memory and Cognition, 43(12), 1962–1987. 10.1037/xlm0000421

120. Vrieze, E., Pizzagalli, D. A., Demyttenaere, K., Hompes, T., Sienaert, P., de Boer, P., Schmidt, M., & Claes, S. (2013). Reduced Reward Learning Predicts Outcome in Major Depressive Disorder. Biological Psychiatry, 73(7), 639–645. 10.1016/j.biopsych.2012.10.014

121. Wager, T. D., & Smith, E. E. (2003). Neuroimaging studies of working memory: A meta-analysis. *Cognitive*, Affective & Behavioral Neuroscience, 3(4), 255–274. 10.3758/CABN.3.4.255

122. Wagner, F., Schmuki, R., Wagner, T., & Wolstenholme, P. (2006). Modeling Software with Finite State Machines — A Practical Approach. Auerbach Publications.

123. Walker, S. C., Robbins, T. W., & Roberts, A. C. (2009). Response Disengagement on a Spatial Self-Ordered Sequencing Task: Effects of Regionally Selective Excitotoxic Lesions and Serotonin Depletion within the Prefrontal Cortex. Journal of Neuroscience, 29(18), 6033–6041. 10.1523/JNEUROSCI.0312-09.2009

124. Wardak, C., Ibos, G., Duhamel, J. R., & Olivier, E. (2006). Contribution of the monkey frontal eye field to covert visual attention. Journal of Neuroscience, 26(16), 4228–4235. 10.1523/JNEUROSCI.3336-05.2006

125. Watson, M. R., Voloh, B., Naghizadeh, M., & Womelsdorf, T. (2019a). Quaddles: A multidimensional 3-D object set with parametrically controlled and customizable features. Behavior Research Methods, 51, 2522–2532. 10.3758/s13428-018-1097-5

126. Watson, M. R., Voloh, B., Thomas, C., Hasan, A., & Womelsdorf, T. (2019b). USE: An integrative suite for temporally-precise psychophysical experiments in virtual environments for human, nonhuman, and artificially intelligent agents. J Neurosci Methods, 326, 108374. 10.1016/j.jneumeth.2019.108374

127. Weed, M. R., Taffe, M. A., Polis, I., Roberts, A. C., Robbins, T. W. A., Koob, G. F., Bloom, F. E., & Gold, L. H. (1999). Performance norms for a rhesus monkey neuropsychological testing battery: acquisition and long-term performance. Cognitive Brain Research, 8(3), 185–201. 10.1016/S0926-6410(99)00020-8

128. Westbrook, A., van den Bosch, R., Maatta, J. I., Hofmans, L., and Papadopetraki, D., Cools, R., & Frank, M. J. (2020). Dopamine promotes cognitive effort by biasing the benefits versus costs of cognitive work. Science, 367(6484, SI), 1362+. 10.1126/science.aaz5891

129. Wolfe, J. M. (1992). Guided search 2.0-A revised model of visual-search. Bulletin of the Psychonomic Society, 30(6), 485.

130. Wolfe, J. M., Cave, K. R., & Franzel, S. I. (1989). Guided search-an alternative to the feature integration model for visual-search. Journal of Experimental Psychology-Human Perception and Performance, 15(3), 419–433. 10.1037/0096-1523.15.3.419

131. Womelsdorf, T., Thomas, C., Neumann, A., Watson, M. R., Banaie Boroujeni, K., Hassani, S. A., Parker, J., & Hoffman, K. L. (2021). A Kiosk Station for the Assessment of Multiple Cognitive Domains and Cognitive Enrichment of Monkeys. Front Behav Neurosci, 15, 721069. 10.3389/fnbeh.2021.721069

132. Wood, S. J., Pantelis, C., Proffitt, T., Phillips, L. J., Stuart, G. W., and Buchanan, J. A., Mahony, K., Brewer, W., Smith, D. J., & McGorry, P. D. (2003). Spatial working memory ability is a marker of risk-for-psychosis. Psychological Medicine, 33(7), 1239–1247. 10.1017/S0033291703008067

